# Improving inversion of model parameters from action potential recordings with kernel methods

**DOI:** 10.1101/2023.03.15.532862

**Authors:** Andreas Oslandsbotn, Alexander Cloninger, Nickolas Forsch

## Abstract

Current methods for solving inverse problems in cardiac electrophysiology are limited by their accuracy, scalability, practicality, or a combination of these. In this proof-of-concept study we demonstrate the feasibility of using kernel methods to solve the inverse problem of estimating the parameters of ionic membrane currents from observations of corresponding action potential (AP) traces. In particular, we consider AP traces generated by a cardiac cell action potential model, which mimics those obtained experimentally in measurable *in vitro* cardiac systems. Using synthetic training data from the 1977 Beeler-Reuter AP model of mammalian ventricular cardiomyocytes, we demonstrate our recently proposed boosted kernel ridge regression (KRR) solver StreaMRAK, which is particularly robust and well-adapted for high-complexity functions. We show that this method is less memory demanding, estimates the model parameters with higher accuracy, and is less exposed to parameter sensitivity issues than existing methods, such as standard KRR solvers and loss-minimization schemes based on nearest-neighbor heuristics.

## 1 Introduction

Measurement of ionic currents in cardiomyocytes can provide key insights into the mechanisms of dysfunctional cardiac electrical properties, with important applications such as cardiac antiarrhythmic drug development. While direct measurement of ionic current modulation has advanced from laborious, manual, low-throughput patch clamp techniques to higher-throughput, multi-cell platforms, these platforms require highly specialized laboratory expertise and high initial expense for the instrumentation [1, 2].

Meanwhile, the dynamic cardiomyocyte membrane potential, commonly referred to as the action potential (AP), is accessible optically using live cell fluorescence microscopy and more directly via microelectrode arrays for certain *in vitro* model systems [3, 4, 5]. The AP is usually recorded over some time interval, and the corresponding time series is commonly called an AP trace. The AP trace and its characteristic shape are determined by the biophysical dynamics of these ionic membrane currents [6, 7]. Furthermore, numerous cardiac action potential models are developed to describe the relationship between the AP and the underlying ionic currents [8, 9, 10, 11, 12, 13, 14, 15, 16, 17]. This makes it possible, at least in theory, to quantify ionic currents from observations of the AP through the use of an AP model. The challenge is that existing AP models are constructed to describe the mapping from ionic currents to the AP, and we are interested in the inverse relationship; namely, given an AP trace, we want the underlying currents.

We can formulate this problem mathematically by representing the ionic membrane currents as a vector *p* = (*p_i_, … p_d_*) ∈ *P* ⊆ *R^d^* and the corresponding AP trace as generated by an unknown function *v* = *f* (*p*). AP models construct a function *f* that approximates *f*, typically described as a system of ODEs or PDEs. The task of deriving ionic currents from AP trace measurements can then be defined as the inverse problem in Definition 1.1.

**Definition 1.1** (The inverse problem). Given an AP trace w find the underlying ionic membrane currents p_w_ = *f^−^*^1^(*w*) *using the AP model f* .

The usual approach in the literature for solving this inverse problem is to define a loss function *L* on a set of AP features {*ϕ_i_*(*v*)}*^m^* (represented as a vector *ϕ*(*v*) = (*ϕ*(*v*)_1_*, …, ϕ*(*v*)*_m_*) ∈ R*^m^*) and seek to minimize this loss function using numerical optimization schemes. The AP features are designed to capture important biophysical properties of the AP traces, and their construction generally requires specialist domain knowledge. Commonly used optimization schemes are direct search methods (nongradient-based), such as Particle swarm [18], Pattern-search, Nelder-Mead, genetic algorithms [19, 20] or combinations of these methods [21]. For example, Jæger et al. [22] used Nelder-Mead combined with the continuation method [23]. The strategy of these methods is to search iteratively in the parameter space P, and for each candidate, *p* ∈ P, solve the ODE model *f* (*p*) to compare with the target *w* through the loss *L*(*ϕ*(*w*)*, ϕ*(*f* (*p*))).

The problem with these classical inversion strategies is their low scalability, which makes them inefficient in the face of large quantities of measured APs. This is because only one AP trace can be inverted at a time, and the AP model *f* must be computed for every candidate parameter. Although the former can be alleviated by parallelization, the computational expense of solving the AP model constitutes a computational bottleneck that prevents the scalability of traditional numerical inversion methods for this setting. Moreover, this computational bottleneck is growing since cardiac electrophysiology models are becoming increasingly large and elaborate [24, 25] and, therefore, more expensive to compute. In addition to this, iterative optimization schemes are often vulnerable to local minima. This is problematic, especially in clinical applications where high reliability is essential, as wrong predictions can have severe consequences.

In addition to issues with scalability and reliability, the inversion of AP traces in cardiac electrophysiology faces the issue of parameter identifiability. This issue typically occurs when more than one parameter affects the AP so that their joint effect is significantly reduced or canceled [26]. This is especially problematic in large complex models, which include multiple inward and outward currents. A special case of this issue is when a parameter has low sensitivity, namely a small (hard to detect) effect on the AP [27]. The issue of parameter identifiability has been studied in several works [28, 26, 29, 27]

In general, successful AP inversion schemes should be able to satisfactorily address the following aspects:

1. *Scalability*, both in the sense of the number of parameters to estimate and in the sense of the quantity of data to parameterize.
2. *Reliability* over the parameter domain, both in the sense of accuracy and in the sense of variability in accuracy over the parameter domain
3. *Identifiability* and *sensitivity* issues with model parameters.

a. Sensitivity issues: Due to parameters that have a small (hard to detect) effect on the AP trace (such as the fast inward sodium (Na+) current in the 1977-Beeler-Reuter model)
b. Identifiability issues: Due to parameters whose corresponding effects (i.e., currents) are overlapping such that the combined effect on the action potential is hard to separate into two or more currents.
4. *Other considerations*: Analysis that is specific to studying AP dynamics under the effects of various drug compounds, such as cycle length and sampling rate.

Tveito et al. [30] introduced an alternative to traditional inversion strategies by optimizing over a pre-computed database of samples 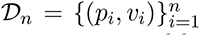 using nearest neighbor heuristics [30]. The advantage of this approach is that the computational burden is moved from the prediction step to a pre-computing phase, and the prediction of parameters for different AP can reuse the same database. Consequently, the algorithm is significantly more scalable when estimating the parameters of several measured AP traces. The challenge with this approach is the large memory requirements of storing the data set. Furthermore, the accuracy is directly linked to the density of the pre-computed samples, a problem not present in the inversion schemes mentioned before. In particular, the accuracy of out-of-sample parameter estimations can only be as good as the distance to the nearest parameter in the pre-computed samples *D_n_*. In other words, reliability would be expected to vary across the parameter domain as a function of the distance to samples in *D_n_*.

In this study, we offer a different strategy, namely, to learn a model of the inverse map *f^−^*^1^ (the mapping from AP traces to the underlying ionic membrane currents). To achieve this, we propose to use the recently developed streaming multi-resolution adaptive kernel algorithm StreaMRAK [31], a kernel method based on kernel ridge regression (KRR). This is a natural choice since kernel methods such as StreaMRAK and other state-of-the-art KRR solvers such as FALKON [32] have already been proven as large-scale learning schemes [32, 33, 31].

Furthermore, kernel methods can generate highly non-linear features of the AP trace in a data-driven manner that can replace the tailored AP features used in traditional methods. The feature space generated by kernel methods allows the modeling of highly non-linear functions without the need for explicit assumptions on the shape and structure of the function; this gives promise for modeling highly involved relationships between AP traces and ionic currents. Moreover, KRR is a convex optimization problem, which rules out local minima.

The goal of this proof-of-concept study is to introduce kernel methods as a new approach to the inverse problem in cardiac cell parameter estimation. The focus is, in particular, to address the issues of scalability (1) and reliability (2) exhibited by existing methods. We also study how our new method performs with respect to the aforementioned parameter sensitivity issues (3a), i.e., parameters whose effect on the AP is difficult to detect. Analysis of identifiability issues due to currents that compound or negate each other will remain to be analyzed in a future study. Similarly, will other issues related to specific AP dynamics.

For a thorough analysis, the study is restricted to a selected subset of parameters in a well-studied model of cardiac cell electrophysiology, the Beeler-Reuter (1977) model of mammalian ventricular cardiomyocytes [34]. The BR model was the first ventricular cell AP model to be developed and included four different ionic currents: a fast inward sodium (Na^+^) current, a slow inward current carried primarily by calcium (Ca^2+^) ions, a time-dependent outward current, and time-independent potassium (K^+^) current. Many cardiac AP models of various cell types have been developed since the BR model was first introduced, incorporating more detailed electrophysiology through the representation of more specific currents, both in the cellular and sub-cellular membranes and compartments.

Of particular interest is the fast inward sodium (Na^+^) current, which affects the slope and peak of depolarization. This current is well known to be difficult to characterize due to its short activation relative to the entire AP and the low degree of sensitivity of the AP to changes in this current [35]. We will study how the proposed inversion scheme handles a parameter sensitivity problem using the Na^+^current.

To demonstrate our approach, we run our method and others on several synthetic experiments on data generated from the BR model. The contribution of our study is summarized in the following; see also Figure 1 for a comparison to existing methods.

- Our approach relies on a pre-computed database, similar to Tveito et al. [30], which allows better scalability compared to standard inversion schemes.
- By learning a model, instead of relying directly on the dataset, our proposed method reduces memory requirements relative to the scheme used in Tveito et al. [30], from O(*n*) to 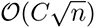, at the cost of an upfront computational training time. Here *C* is a constant independent of *n*.
- We demonstrate with synthetic experiments how StreaMRAK can estimate model parameters with increased reliability compared to alternative methods.
- For the analysis, we propose a geometric approach to identifiability analysis, which supplements the tool developed in Jæger et al. [27].

**Figure 1:**
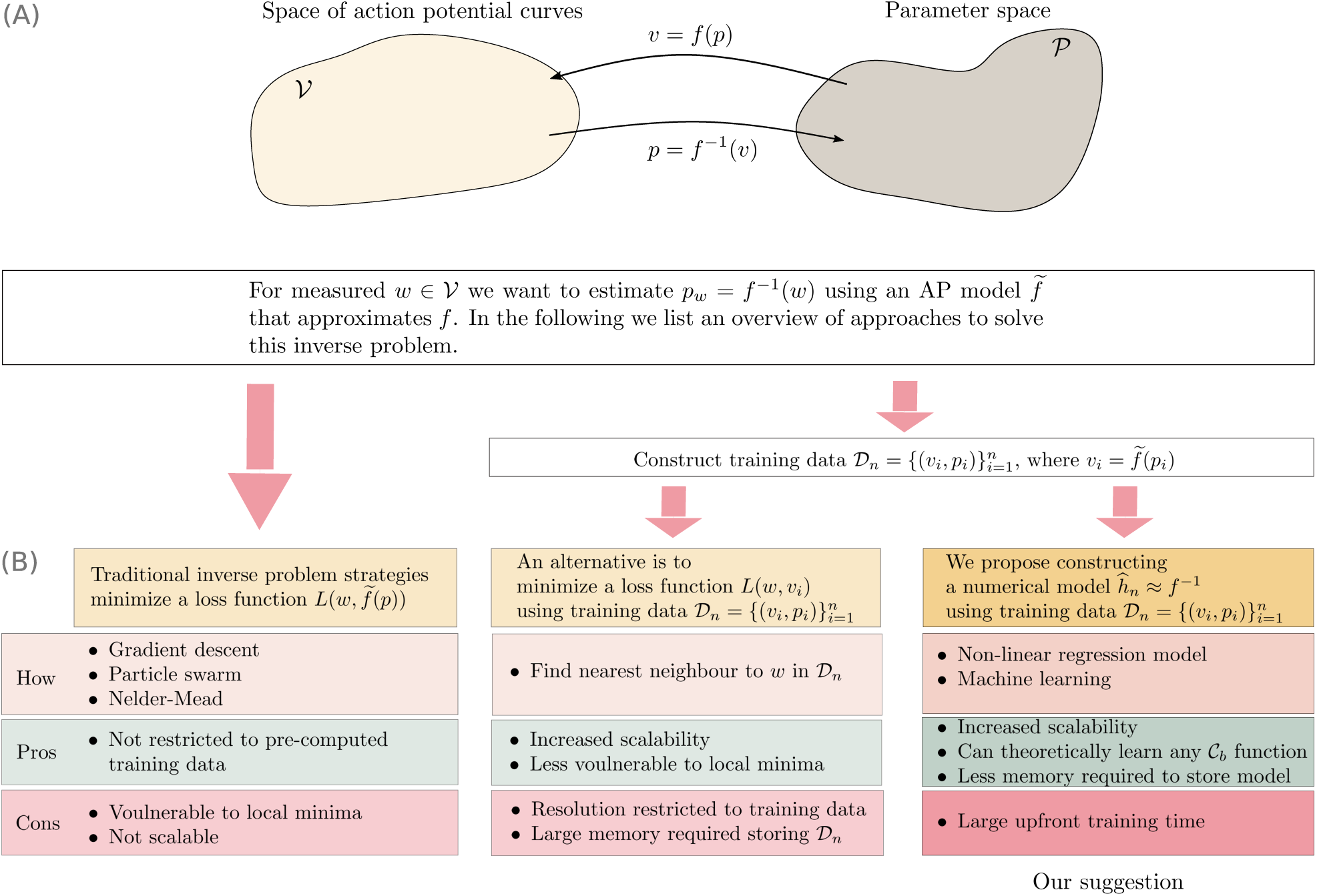
Panel (A) illustrates the mapping *f* and its inverse *f^−^*^1^ between the parameter space 𝒫 and the voltage space V and defines the inverse problem we want to solve. Panel (B) compares our method with existing inverse problem strategies. Here C*_b_* is the set of bounded and continuous functions.

The current study provides a solid foundation with valuable insights for further analysis. In future work, we will focus on extending the analysis to more complex AP models with more parameters, states, and variables, such as Tusscher and Panfilov [9] and O’Hara et al. [11] or even models of cardiomyocytes derived from human induced pluripotent stem cells, such as Paci et al. [36].

## 2 Methods and Theory

In this section, we offer a detailed description of the method we propose for learning the inverse map between AP traces and the underlying parameters. We start by describing how we generate the synthetic training data. In particular, we present the 1977 Beeler-Reuter model, which is the AP model we consider in this study. Section 2.2 describes the kernel model we propose for the non-linear regression and the training scheme used to fit the model. Meanwhile, Section 2.3 briefly recaptures the loss-minimization schemes we compare with, and Section 2.4 offers an outline of specific parameter identifiability tools that we use to analyze the AP model for better understanding of the estimation results. Finally, Section 2.5 gives an overview of the study and the experiments we conduct.

### 2.1 Generating synthetic data from the AP model

The method we propose requires access to a set of training data 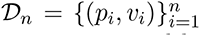, where *p_i_* ∈ R*^d^* is a *d* dimensional vector of parameters representing different ionic membrane currents and *v_i_* is a time series of transmembrane potential (voltage) *v_i_*, which we refer to as the action potential (AP). The training data can be generated either from *in-vitro* measurements or numerically solving an AP model. Although the transmembrane potential is a continuous function of time, in practice, whether we measure the AP traces *in-vitro* or generate them from a system of ODEs, we need to sample the traces at a finite time grid. We denote this time grid [*t*_1_*, …, t_k_, … t_T_*], which we scale such that *t*_1_ = 0 and *t_T_* = 1. We think of these AP traces *v_i_* as *T* dimensional vectors in the sub-space *V_T_* ⊂ *R^T^*, where each entry *v_ij_* corresponds to a time step *t_j_*. See panel (C) in Figure. 2 for an illustration.

**Figure 2:**
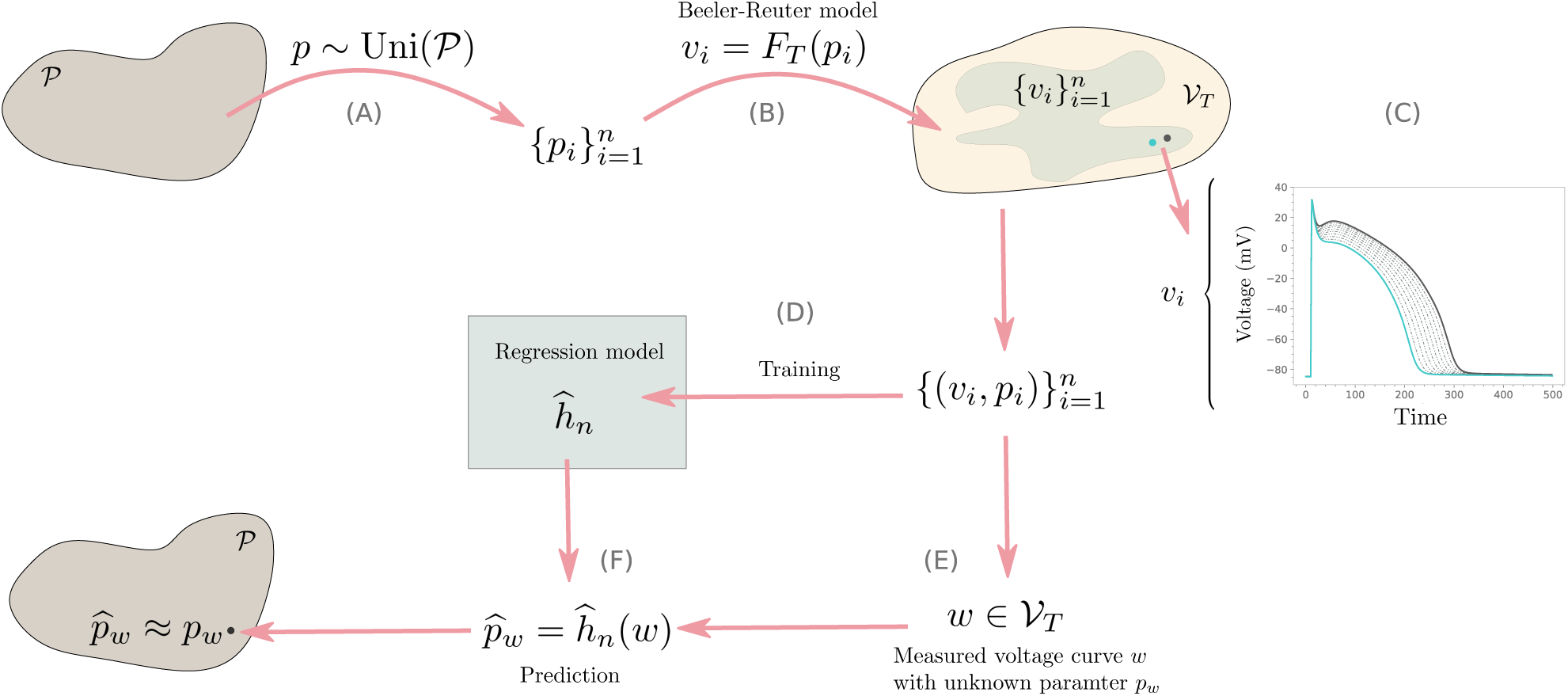
Illustration of our method

Now, training samples generated by an AP model with the parameter set *p* = (*p*_1_*, …, p_d_*) restrict predictions to these parameters. For a given AP model *f*, we consider some physiologically viable region 𝒫 ⊆ R*^d^* from which we sample *n* parameters uniformly, and solve the BR model on each sample, generating the training data 𝒟*_n_* = {(*f* (*p_i_*)*, p_i_*) : *p_i_* ∼ Uni(P), for *i* = 1*, … n*}. See step (A)-(B) in Figure 2. In the remainder of this paper, we use the 1977 Beeler-Reuter AP model of mammalian ventricular cardiomyocytes to generate the training samples. We will now give a brief description of this model.

#### 2.1.1 The action potential model

The Beeler-Reuter model is formulated as a system of ODEs relating the time derivative of the action potential to a set of parameters representing specific membrane currents. In this study, the parameters we consider *p* = (*p*_1_*, … p_d_*) are scaling parameters that alter the amplitude of ionic currents. Table 1 summarizes these parameters and the ionic currents they represent; once solved numerically, the model outputs a time series of transmembrane potential (voltage) *v_i_*, which we refer to as the AP. For a more detailed list of stimulus currents and other variables of the ODE implementation, we refer to Table S.3 in the Supplementary.

**Table 1:** Scaling parameters that alter the amplitude of ionic currents in the Beeler-Reuter model. The first column lists the parameter names. The second contains scaling constants, and the last column contains a description.

| Parameter | Scaling | Description |
| --- | --- | --- |
| $g_{Na}$ | $4.0 \times 10^{-2} \text{ mS/mm}^2$ | Sodium ($\text{Na}^{2+}$) ionic current component ($\text{mS/mm}^2$) |
| $g_s$ | $9.0 \times 10^{-4} \text{ mS/mm}^2$ | Slow inward ionic current component ($\text{mS/mm}^2$) |
| $g_K$ | $3.5 \times 10^{-3} \text{ mS/mm}^2$ | Time-independent potassium ($\text{K}^+$) current ($\text{mS/mm}^2$) |
| $g_{x1}$ | $1.9 \times 10^{-3} \text{ mS/mm}^2$ | Time-dependent outward current ($\text{mS/mm}^2$) |

The ODEs that make up the Beeler-Reuter model are described mathematically in Halfar [37]. However, instead of working directly with the ODE equations of the Beeler-Reuter model, it is sufficient for our purposes to consider the model as a mapping from the parameter space 𝒫 to the space of AP traces 𝒱*_T_*, we defined this mapping in Definition 2.1. See also step (B) in Figure 2 for an illustration on how we use the model.

**Definition 2.1** (The Beeler-Reuter function). Under the setting described above, the Beeler-Reuter model corresponds to a vector-valued function *F_T_* : *P* → *V_T_ . In particular,*

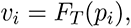

*where v_i_* = (*v_i_*_1_*, …, v_ik_, …, v_iT_*) ∈ V *is the AP curve that corresponds to the parameter choice p_i_* = (*p_i_*_1_*, …, p_ij_, …, p_id_*) ∈ *P . Furthermore, the action potential at time t_k_ is*

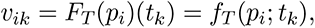

*where each time step t_k_ defines a specific functional relationship between the parameters and the AP*.

#### 2.1.2 Pre-processing of the AP traces

We use single pulse AP traces *v_i_* paced at 1 Hz with a sampling rate of 1 kHz, where only the first half of the pulse is included for the regression. Because the waveform is contained in the first 500 ms of the AP pulse, this reduces the problem’s dimensionality without losing any information about the shape of the waveform. The time interval is chosen to be sufficiently large for all variations of AP traces over the domain 𝒫 to re-polarize to the resting state.

Furthermore, the AP recording methods we are trying to simulate with our synthetic data do not obtain absolute measurements of the resting potential. Because of this, we align the resting potential for all traces in our experiments to represent the likely case where we do not have this information.

After these pre-processing steps, we consider the resulting single-pulse AP traces *v_i_*as *T* dimensional vectors in the sub-space *V* ⊂ R*^T^* .

#### 2.1.3 Pre-processing of the parameters

The parameters of the Beeler-Reuter model are of different units and magnitudes. For a stable numerical analysis, we scale the parameters to unitless quantities using the scaling constants in the second column in Table 1. Under this scaling, the parameter *p* = (1*, …,* 1) corresponds to what we define as a baseline biophysical cell with its corresponding AP trace. This way, a 0.1 perturbation of any parameters corresponds to a 10% change in their magnitude, etc.

#### 2.1.4 Choice of parameters for demonstrating parameter estimation capabilities

The goal of this work is to study the parameter estimation capability of our method compared to existing schemes with the purpose of creating a solid foundation for further application of the proposed method to more complex models. With this intent, we have chosen to use the fast inward Na^+^ current and the slow inward Ca^2+^-type current to demonstrate how the method can capture distinctly different changes to the AP trace with high reliability.

We note that the currents being studied here are commonly examined when investigating anti- and pro-arrhythmic effects of cardiac drugs [35] and are, therefore, natural choices in their own right. Furthermore, modulations of the fast Na^+^ current are known to be difficult to detect due to it only being active during the fast upstroke of depolarization and considering the sampling constraints of modern measurement systems. Estimating the Na^+^ current is, therefore, an interesting challenge for the inversion model when evaluating precision and reliability. Furthermore, the analysis is constrained to two model parameters to promote visualization and interpretation of the analysis. The authors refer the reader to Section 4.4 and 4.5 for more rationale on this choice.

### 2.2 Constructing the kernel models

Having generated the training data 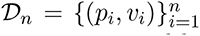 as described in the previous section, we are in a position to construct our non-linear regression model. This modeling strategy corresponds to the rightmost column in panel (B) in Figure 1. We note that the modeling method proposed here is not restricted to training data generated from the Beeler-Reuter model. The training data 𝒟*_n_* might also come from solving an alternative action potential model or from experimental data measured in the lab.

In this section, we outline the main principles of kernel methods as the underlying regression model we use. We then offer a brief sketch of the specific kernel model we propose, namely StreaMRAK [31], and an alternative kernel algorithm FALKON [32], which we compare with. For more details on these kernel methods, we refer to Appendix 6.5 and the references therein.

#### 2.2.1 A note on the inverse map

We begin by making a short remark on the inverse map we set out to model. We are interested in constructing a non-linear regression model that can approximate the inverse map *f^−^*^1^, illustrated in panel (A) in Figure 1. Since our training data is generated by the Beeler-Reuter model *v* = *F_T_* (*p*), we are, in practice, building a model that approximates *p* = *F^−^*^1^(*v*). In Appendix 6.3, we discuss the invertibility of this mapping in more detail. Furthermore, each of the parameters we consider *p_j_, j* ∈ 1*, …, d* are independent (orthogonal), which means they can be described by independent mappings from the voltage space. We represent each of these mappings as

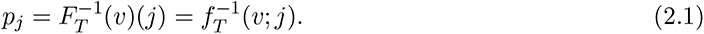

#### 2.2.2 Defining the model

Appendix 6.5 gives a more detailed description of the kernel model. Here we restrict ourselves to presenting the main building blocks. The model we propose to approximate *f^−^*^1^(*v*; *j*) corresponds to a linear combination of kernels *k*(*v, v_i_*) centered on the training data:

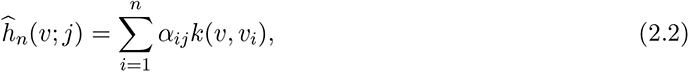

where our choice of *k*(*v, v_i_*) is the Gaussian kernel

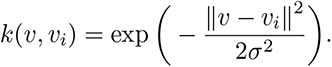

The advantage of this model is that *k* is a universal kernel [38], meaning that *h_n_* can approximate bounded and continuous functions on a domain X arbitrarily well provided sufficient training data.

Furthermore, since the model is non-parametric, we can capture highly complex functions without any background knowledge of the functional characteristics. StreaMRAK introduces a multi-resolution formulation of the model in (2.2). Details on this are given in Appendix 6.5.

We note that with the choice of the Gaussian kernel, the model used in Eq. (2.2) corresponds to interpolation with radial basis functions. Interpolation with radial basis functions is a common choice in machine learning and regression, for which the task is to learn the weights of the expansion. Numerous approaches exist for learning these coefficients, focusing on efficiency and the bias-variance tradeoff; an overview of different approaches can be found in [39]. The approach we use for this purpose is based on kernel regularized ridge regression (KRR) which is supported by a rich theoretical foundation that guarantees the method’s reliability in terms of bounds on the bias and variance of the approximation. In the next section, we discuss the methods we use for finding the coefficients in more detail.

#### 2.2.3 Learning the coefficients

Training the kernel model from Eq. (2.2) corresponds to finding an optimal set of regression coefficients *α_ij_*. We will use two recently proposed algorithms for this purpose: The large-scale kernel method FALKON [32] and the multi-resolution adaptive kernel method StreaMRAK [31]. FALKON is rooted in the optimization scheme known as kernel ridge regression (KRR), see Eq. (6.1) in the appendix, but introduces several improvements to speed up computations. Meanwhile, StreaMRAK utilizes FALKON as a base solver but introduces a boosted version to construct a model that can learn highly complex functions with varying modalities; large local derivatives in some regions and derivatives near zero in others.

We mention that both FALKON and StreaMRAK use sub-sampling of the training data to reduce computational complexity. In FALKON, random sub-sampling, also known as Nystrom sub-sampling, is used to select a subset of training points on which the regression is performed. Meanwhile, StreaMRAK uses a cover tree to construct an epsilon cover in the AP space V. The epsilon cover enforces a minimal distance of *ϵ* between each AP trace *v_i_* ∈ V and also tries to maintain a maximal distance of *ϵ*. This has the advantage of removing training data that contribute with similar information (training points that are very close) but simultaneously ensuring that training points are close enough to maintain sufficient interpolation properties.

#### 2.2.4 Parameter estimation

The trained model *h_n_* can be used to estimate the underlying parameters of measured AP traces *w* ∈ *V_T_* for which the corresponding set of parameters *p_w_* is unknown. These AP traces can be, for example, time series measurements of the cardiac transmembrane potential using live cell fluorescence microscopy, as described in [4]. In this regard, the time grid underlying the training data must be the same as for the AP traces *w* measured *in-vitro*. If this is not the case, then the AP traces are not comparable. In other words, we must ensure the measured AP traces *w* belong to the same space *V_T_* as the training data, by embedding them in the same vector space.

However, should the time grids differ initially, it is possible to align them by time interpolation or down sampling, which is possible because the AP traces are smooth functions of time. This way, one can avoid training the model again for different experimental recording systems that differ in sampling frequency.

Provided *w* ∈ *V_T_*, we can make estimations of the corresponding parameters using the model *ĥ_n_*, such that *p_w_* = *ĥ_n_*(*w*). This is illustrated in step (E)-(F) in Figure 2. We note that for our synthetic experiments, we do not use AP traces measured experimentally. Instead, we generate a separate out-of-sample test set 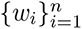 using the BR model on which we demonstrate our approach.

### 2.3 The nearest neighbor loss-minimization inversion schemes

For comparison with the regression strategy, we implement two variations of the nearest neighbor loss function minimization strategy; see the middle column in panel (B) in Figure 1. Given a measured AP trace *w* ∈ 𝒱*_T_* and a pre-computed training set 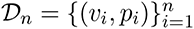 the methods finds

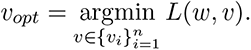

Essentially, this corresponds to finding the nearest neighbor of *w* among the *v_i_* ∈ 𝒟*_n_*, when ”distance” is measured using *L*.

We consider two loss functions for finding the nearest neighbor. The simplest loss function is 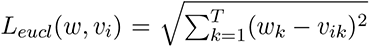, which simply measures the euclidean distance between AP traces in 𝒱*_T_* . In addition, we use a loss function *L_apf_* (*w, v_i_*) = *_j_ H_j_*(*w, v_i_*)^2^ that measures the distance using specific action potential features (APF) *ϕ_j_*, such as AP duration. A complete list of APF and detailed descriptions can be found in S3.1 of Jæger et al. [22].

The motivation for finding *v_opt_* is that this provides a lower bound on the prediction accuracy using the bounding box search used in Tveito et al. [30]. Clearly, for sufficiently large *n*, finding *v_opt_* is computationally impractical, which is also why Tveito et al. [30] instead made use of an iterative search in bounding boxes (subsets) of 𝒟*_n_*. However, for our purposes, we are interested in accuracy and reliability comparisons, and as such, *v_opt_* is a natural choice.

### 2.4 Identifiability analysis

We are interested in learning the relationship between action potential traces and the parameters of the underlying ionic currents. However, as noted in Jæger et al. [27], typical AP models have parameters that result in similar or overlapping changes, and the sensitivity of the AP to parameter perturbations can vary considerably. Because of this, we expect that the effect of some parameters is more challenging to learn than others. By knowing which directions are expected to be hard to learn, one can account for that in the design of the learning model. Furthermore, knowing the parameter identifiability is also valuable when assessing the reliability of predictions.

In this section, we review an existing strategy, proposed in Jæger et al. [27], for determining parameter sensitivity and identifiability based on spectral analysis of a matrix that incorporates the currents and their time dependency. Furthermore, we suggest a supplementary tool based on the Laplacian eigenmaps method [40], which offers a geometric understanding of the mapping between parameters and AP traces.

#### 2.4.1 Spectral analysis

Jæger et al. [27] proposed a tool for analyzing the identifiability of ionic currents in a given AP model using spectral analysis. Let *I_j_^k^* be membrane current *j* at time step *t_k_* = *k*Δ*t* and form the matrix

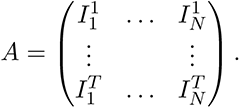

The total membrane current can then be written as *I_tot_* = *Aµ*, where *µ* ∈ ℝ*^N^* is the vector of only ones.

Under the assumption that the AP is determined by *I_tot_*, the identifiability of each current component, *I_j_*, can be determined by considering the singular value decomposition of A. In particular, *I_tot_* and therefore the AP is only affected by the currents *I_J_* that project onto the singular vectors associated with non-zero singular values. Because of this, any current in the null space of *A* can be considered non-identifiable.

Furthermore, the change in AP when perturbing along a singular vector is proportional to the corresponding singular value. Consequently, currents projecting along singular vectors with small singular values have less effect on the AP, and changes in these currents are, therefore, harder to detect. Because of this, the size of singular values can be used to understand the identifiability of the membrane currents incorporated in an AP model. We note that the identifiability properties of different AP models vary, as seen in Jæger et al. [27, 22].

#### 2.4.2 Geometrical analysis

We propose to use Laplacian eigenmaps (LE) [40, 41] as a geometrical tool to probe the relationship between AP traces and parameters. The LE is a well-established method for interrogating the structure of non-linear and complex point clouds and is supported by a solid theoretical foundation. In particular, LE can be considered a non-linear counterpart of principal component analysis.

We can think of the set of AP traces {*v_i_*}*_i_^n^* ∈ 𝒱*_T_* as a point cloud in 𝒱*_T_*, generated from a set of uniformly sampled parameters *p_i_* ∈ Ω from some parameter domain Ω ∈ P. The main principle of LE is to construct the graph Laplacian *L* on {*v_i_*}*_i_^n^*, which then incorporates the structure of the point cloud. To gain access to this structure, it is sufficient to solve for the singular vectors {*u_i_*}*_i_^n^* ∈ *R^n^* and singular values *σ_i_* of *L*. In particular, with *σ*_1_ ≤ *…,* ≤ *σ_n_*, we have that *u*_1_ corresponds to the direction of the most extensive spread. Furthermore, with *r* being the rank of *L*, only the *r* first singular vectors are relevant. For a detailed description of how to construct *L*, we refer to Belkin and Niyogi [40].

The similarity to the spectral analysis proposed in Jæger et al. [27] is apparent, with the difference that we now consider the spectral components of the graph Laplacian *L* instead of the current matrix *A*. Furthermore, using the *r* first singular vectors {*u_i_*}*_i_*_=1_*^r^* we can generate an *r*-dimensional embedding of {*v_i_*}_i_*^n^*, where the singular vectors act as a new set of coordinates for the AP traces.

The AP traces *v_i_* ∈ R*^T^* are encoded as *T* dimensional vectors, where *T* is the number of time steps. Meanwhile, the rank *r* of *L* is directly related to the number of independent parameters that generated {*v_i_*}_i_*^n^*. Although an AP model considers a significant amount of independent parameters, the number of independent parameters is typically significantly less than *T*, which means *r* ≪ *T* . More importantly, restricting the parameters space to only a few parameters, *r* ≤ 3, allows us to visualize the embedding.

### 2.5 Overview of study

In this study, we compare the parameter estimation capabilities of the proposed kernel model StreaMRAK with four loss-minimization schemes similar to those used in Tveito et al. [30] and Jæger et al. [22]. In particular, we consider two loss functions: the standard Euclidean norm in the voltage space and a loss function constructed from action potential features (APF); see Section 2.3 for more details. For each loss function, we do a 1-nearest-neighbor search (Eucl-1-nn and Apf-1-nn) and a 10-nearest-neighbor search (Eucl-10-nn and Apf-10-nn). Furthermore, we also include the kernel model FALKON [32] in the comparison.

In the next section, we define the measures used in this study to compare the different inversion schemes. The subsequent sections describe the synthetic training data and the experiments we conduct.

#### 2.5.1 Measures of reliability

When comparing the reliability of the inversion schemes, we consider the accuracy and how it varies over physiologically relevant regions of the parameter space. This is because the application of the inversion scheme we are demonstrating is ultimately a tool for estimating drug effects from *in vitro* AP recordings. A comparison of the inversion schemes is made in light of what is desirable for drug effect estimation. In particular, an inversion scheme that provides high accuracy on average but has large variability over the parameter domain is undesirable because failure to estimate the effect of a particular drug dose can have negative implications.

Let A ⊂ 𝒫 be such a physiologically relevant region in the parameter space. We then quantify the accuracy as the root mean square error (RMSE) of the parameter estimations in A. Similarly, we quantify the variability in the estimate by the standard deviation (Std) and the maximum error over A.

#### 2.5.2 Training data

Using the 1977 Beeler-Reuter AP model of mammalian ventricular cardiomyocytes, we generate a synthetic data set of *N* = 6000 sample pairs 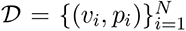, on which we train our models. The source code used for this purpose is taken from Finsberg [42]. The training samples are distributed uniformly over the parameter domain 𝒫 = [0.2, 2]^2^ where the first parameter is the conductance of the rapid and excitatory inward sodium ionic current, *g_Na_*, and the second parameter is the conductance of the slow inward ionic current, *g_s_*, which primarily consists of calcium ions. Furthermore, we assume that *p*_(0)_ = (*g_Na,_*_0_*, g_s,_*_0_) = (1.0, 1.0) are the parameters of the untreated cell, which represents a baseline biophysical cell and standard cardiac action potential.

#### 2.5.3 Outline of experiments

The experiments performed in this study were designed to rigorously compare the accuracy and variability of parameter estimation for the different methods over physiologically relevant regions.

The first experiment, covered in Section 3.1, gives an overview by inspecting the heterogeneity of the estimation errors over the parameter domain P. Meanwhile, the two succeeding sections, Section 3.2 and 3.3, present experiments that inspect the accuracy as a function of specific directions (in P) and distances away from the baseline biophysical cell at *p*_(0)_.

Finally, Section 3.4 demonstrates the proposed inversion scheme as a tool for estimating drug effects using an example with simulated drugs. For this purpose, consider four hypothetical drugs [*A, B, C, D*] whose perturbation directions *θ* are indicated in the right panel of Figure 3. For each drug, we consider a series of parameters *p_θ,r_* = (*θ, r*) for varying radii *r*. From the root mean square and standard deviation over these parameter estimations, we quantify the expected accuracy and variability in the drug-effect estimation for each simulated drug.

**Figure 3:**
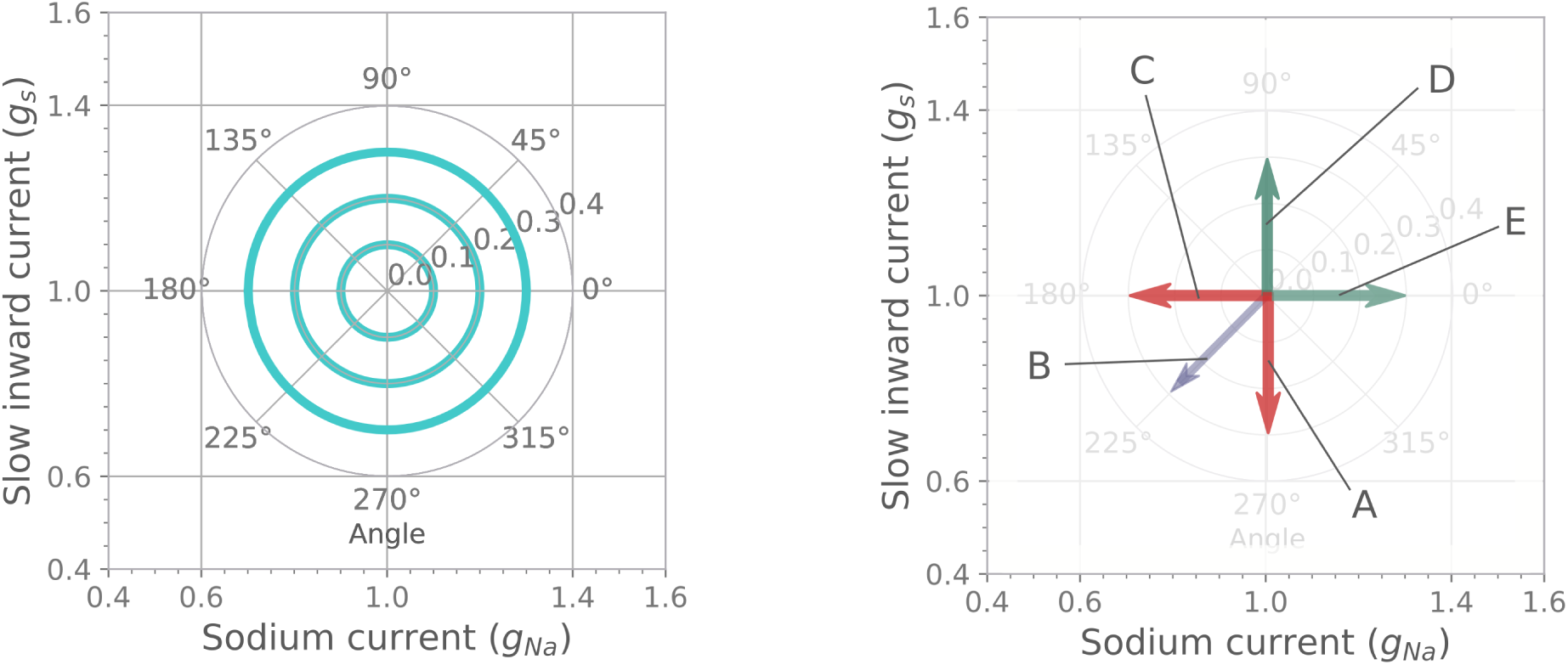
Direction and magnitude of perturbations in parameter space for simulated drug effects on the standard cardiac action potential. The left panel shows the three perturbation magnitudes *r* = 0.1, 0.2, 0.3 for directions uniformly distributed on [0*^◦^,* 360*^◦^*]. The right panel indicates the drug effect of 5 different drugs; A and C are representatives of Ca^2+^ and Na^+^ current inhibitors, respectively; similarly, D and E are representatives of Ca^2+^ and Na^+^ current agonists, and B is a mix of a Ca^2+^ and a Na^+^ current inhibitors.

The experiment conducted in Section 3.2 estimates the parameters of the AP traces along each of the concentric circles in the left panel of Figure 3. The purpose is to examine how the estimation accuracy varies with the direction away from *p*_(0)_ in the parameter space. This experiment allows us to see how the different inversion schemes compare when more than one parameter is perturbed simultaneously.

Meanwhile, the experiment conducted in Section 3.3 examines how the accuracy varies with the distance from *p*_(0)_ along the specific directions in parameter space. In particular, we consider the directions A, B, and C, as illustrated in the right panel in Figure 3, which can be related to the action of relevant drugs acting on the Na^+^- and Ca^2+^-type currents. The purpose is to compare the reliability of the inversion schemes, both in terms of estimation accuracy and consistency, along relevant physiological regions of the parameter space.

We finish the study with a performance analysis in Section 3.5, where we consider the time complexity and parameter estimation capability as a function of model size. An implementation of StreaMRAK and FALKON, along with experiments that reproduce the performance experiments, can be found in the GitHub repository [43].

## 3 Results

This section compares the parameter estimation capabilities of StreaMRAK with FALKON and the loss-minimization schemes. We consider the experiments outlined in Section 2.5.3 and finish the section with a performance analysis on time complexity and parameter estimation capability.

### 3.1 Experiment one: Domain error

This experiment studies the heterogeneity of the estimation error over the parameter domain = [0.2, 2]^2^ to quantify the reliability of parameter estimations from StreaMRAK with respect to the other methods. The reliability over a region is related to both the absolute accuracy of the estimate and how the estimation accuracy varies within this region, with higher variations meaning lower reliability. To generate the test data for this experiment, 3000 points are sampled uniformly from P. The corresponding voltage curves are then solved for using the Beeler-Reuter model. For each test trace, *w_i_*, the underlying parameter vector *p_wi_* ∈ 𝒫 is estimated and compared with the actual parameter *p_i_* from the test set. In this manner, the square error is calculated for each sample in the test data; the distribution of errors over 𝒫 are shown in Figure 4.

**Figure 4:**
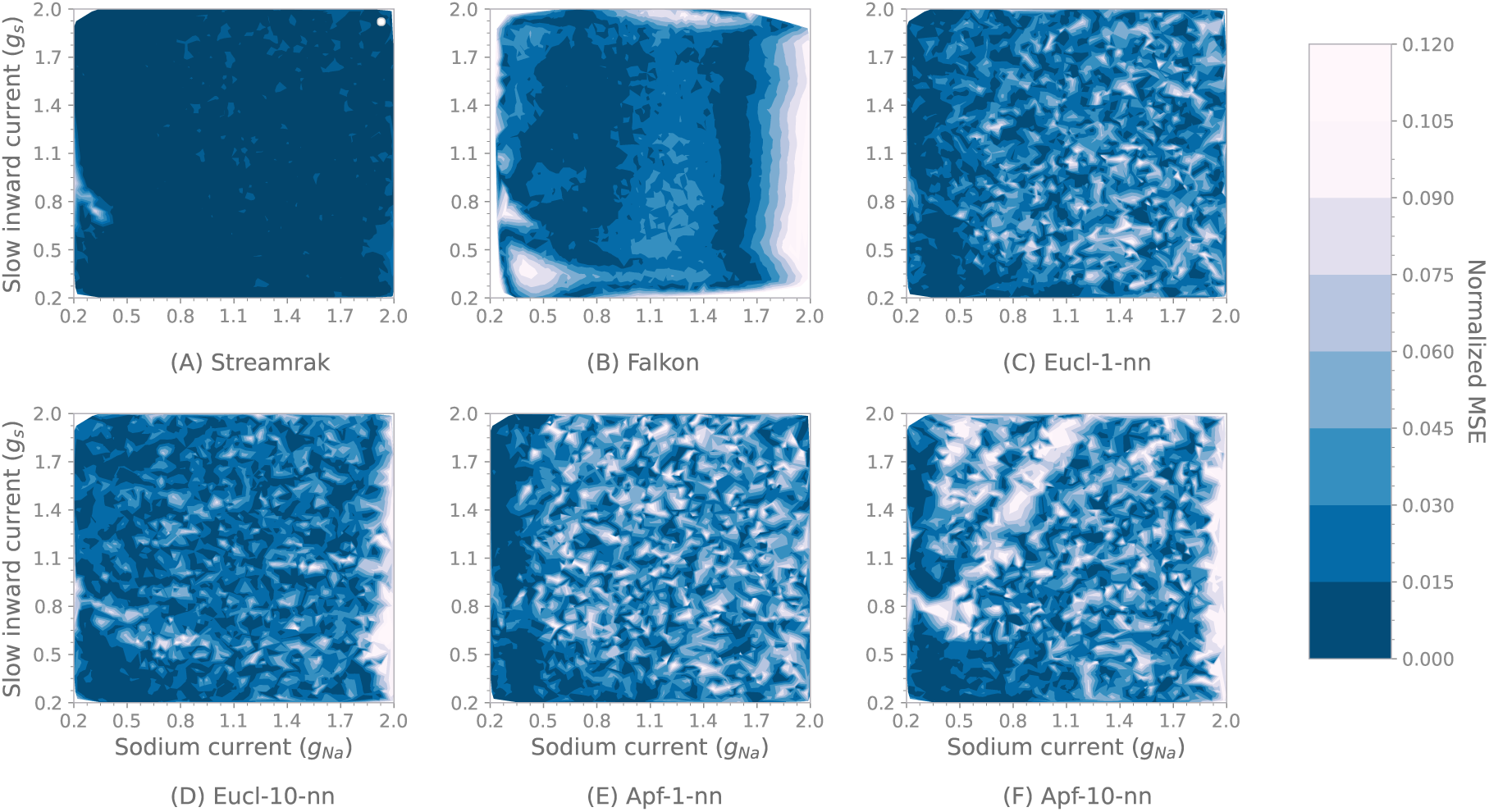
Experiment from Section 3.1. Distribution of estimation error over the domain *P* = [0.2, 2]^2^.

Panel (A)-(B) in figure 4 shows the domain error of the StreaMRAK and FALKON estimations. Both StreaMRAK and FALKON are based on training a non-linear regression model, which interpolates between the samples in the training data D. However, the estimation errors of StreaMRAK are more consistently low compared to FALKON. In particular, FALKON has higher variability and more significant MSE along the domain borders than StreaMRAK.

Panel (C)-(F) shows the domain error of the four loss-minimization algorithms Eucl-1-nn, Eucl-10-nn, Apf-1-nn, and Apf-10-nn. The domain errors of the APF-loss and Euclidean-loss-based methods are relatively comparable; the domain errors of the APF-based methods are slightly larger in magnitude than those based on the euclidean loss function.

By comparing the domain error of the kernel methods (A)-(B) with that of the loss-minimization schemes (C)-(F), one can see the difference between building a regression model of the data, which extends estimations beyond the training samples in 𝒟*_n_*, and minimizing a loss, which is restricted to samples in 𝒟*_n_*. The model-based schemes StreaMRAK and FALKON have significantly more consistent estimations over the parameter domain P. Furthermore, the domain error of StreaMRAK is also considerably lower, whereas, for FALKON, this is only true in some local regions of the parameter domain.

### 3.2 Experiment two: Accuracy as a function of perturbation direction

This experiment studies how the accuracy of the inversion methods varies as a function of direction in parameter space relative to the baseline biophysical cell at *p*_(0)_.

To generate the test data, parameters are sampled uniformly along three concentric circles centered at *p*_(0)_ = (1.0, 1.0) with radius 0.1, 0.2, and 0.3, respectively. For each set of parameters (*g_Na,i_, g_s,i_*) along these perturbation circles, the corresponding AP trace *v_i_* is generated using the Beeler-Reuter model. The underlying parameters of these AP traces are then estimated using each of the six algorithms. By assuming that *p*_(0)_ corresponds to the untreated case, the three circles can be considered as drug perturbations of 10%, 20%, and 30% in all directions in the parameter domain as shown in the left panel in Figure 3.

The parameter estimates from the six algorithms are shown in Panel (A)-(C) in Figure 5 for each of the three perturbation magnitudes, respectively. The accuracy of the parameter estimation from the two kernel methods, StreaMRAK and FALKON, is consistently high throughout the perturbation directions and magnitudes. Meanwhile, the accuracy of the estimations from the euclidean-loss-minimization algorithms Eucl-1-nn, Eucl-10-nn is high around the 90*^◦^* and 270*^◦^* axes, which are directions dominated by perturbations of the slow inward current. However, when perturbations of the Na^+^ current are introduced, the parameter estimation accuracy of these methods drops. We observe similar behavior for the APF-loss-minimization algorithms Apf-1-nn and Apf-10-nn; however, note that for these, the regions with the best estimation accuracy are not along the vertical axes.

**Figure 5:**
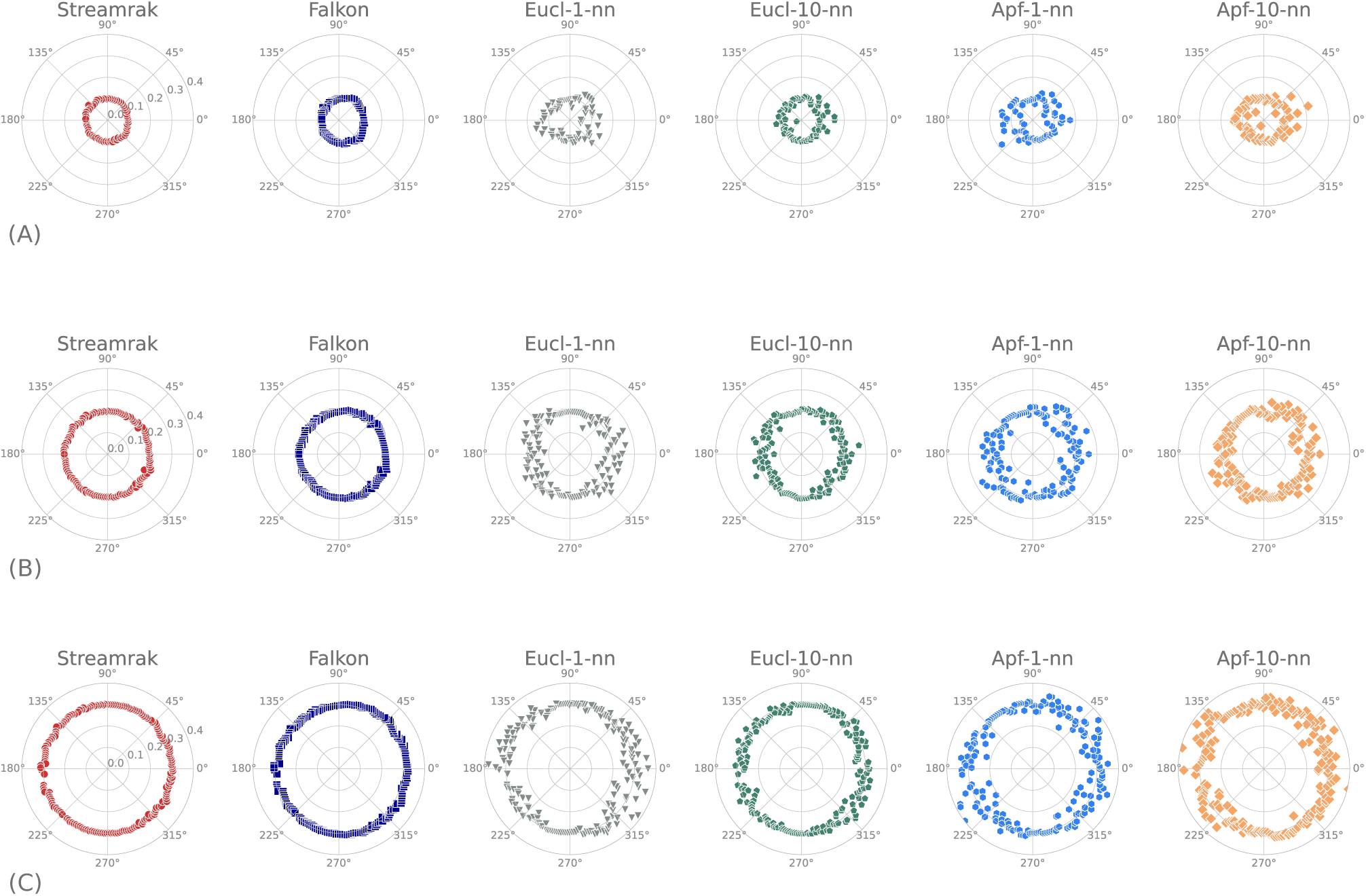
Estimation of perturbations with increasing magnitude for directions in interval [0*^◦^,* 360*^◦^*]. Here panel (A) corresponds to perturbation radius *r* = 0.1 from the unperturbed *p* = (1.0, 1.0) (Smallest perturbation circle in Figure 3). Panel (B) corresponds to perturbation radius *r* = 0.2 (Middle perturbation circle in Figure 3). Panel (C) corresponds to perturbation radius *r* = 0.3 (Largest perturbation circle in Figure 3).

We have observed that the estimation accuracy of the loss-minimization algorithms drops as the magnitude of the perturbation of the Na^+^ current parameter *g_Na_* increases. Furthermore, for the loss-minimization algorithms, perturbations of *g_Na_* affect the estimation accuracy of *g_s_*. Meanwhile, the regression models exhibit more confidence in identifying parameters. Notably, for StreaMRAK and FALKON, the estimation accuracy of *g_s_, g_Na_* does not depend on the perturbation direction and magnitude.

### 3.3 Experiment three: Accuracy as a function of perturbation magnitude

This experiment studies how prediction accuracy varies with perturbation magnitude away from the baseline biophysical cell at *p*_(0)_. For this purpose, the directions A, B, and C introduced in Section 3.4 are considered; See Figure 3. To generate the test data, 20 evenly spaced samples are made on the interval [0, 0.3] along each of the three simulated drug directions. Figure 6 shows the estimation results.

**Figure 6:**
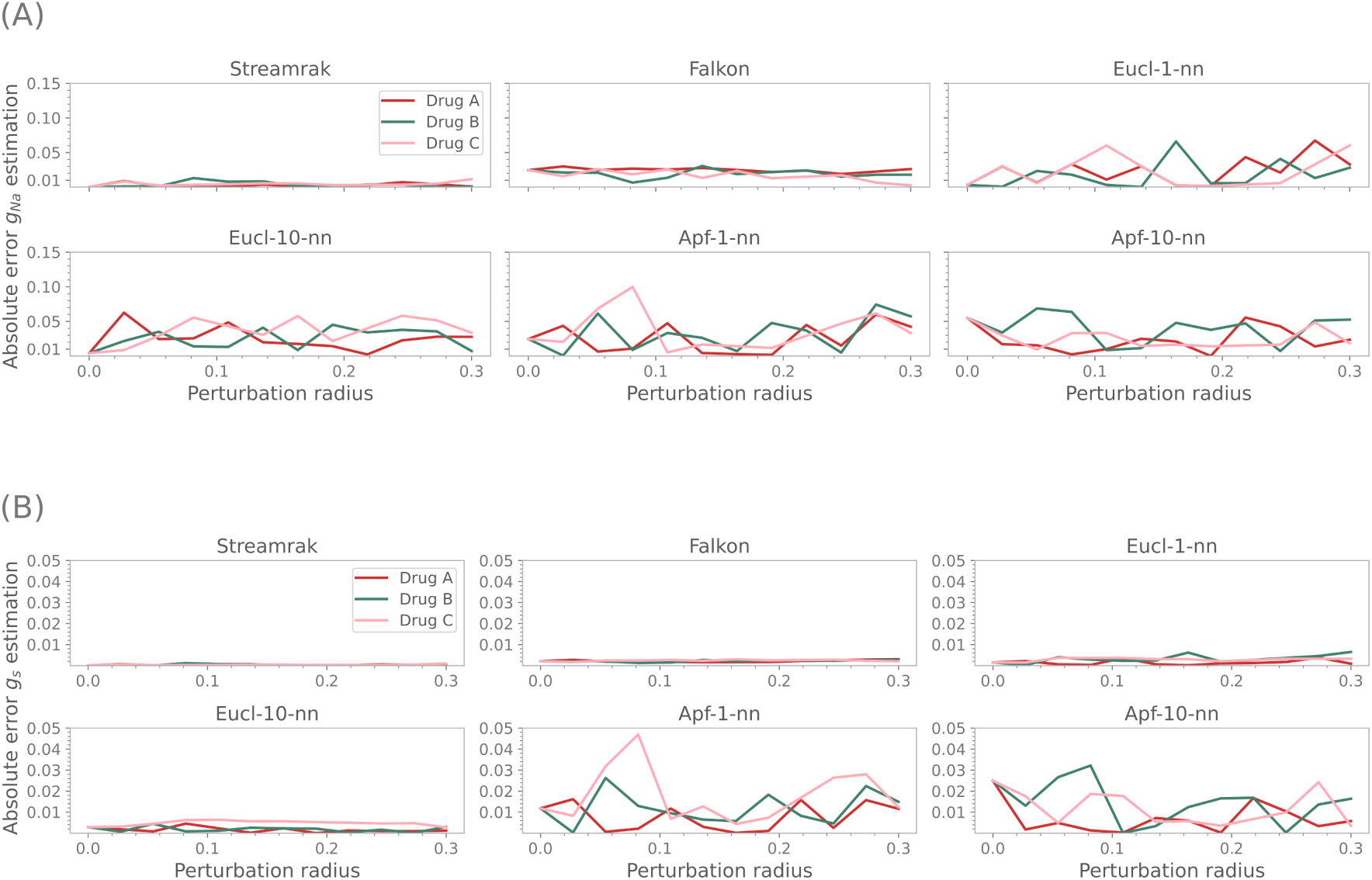
Experiment from Section 3.3. Figure showing parameter estimation accuracy, as measured by the absolute error, along the three selected directions A, B, and C as indicated in Figure 3. For each direction, 20 samples are evenly spaced on the interval [0, 0.3]. The red line is direction A, the green line is B, and the pink line is C. Panel (A) shows the estimation accuracy of the *g_Na_* parameter, and panel (B) shows the estimation accuracy of the *g_s_* parameter.

Panel (A) in Figure 6 shows the estimation error of the *g_Na_* parameter. The parameter estimation accuracy of StreaMRAK and FALKON are significantly more consistent across the perturbation range [0, 0.3] for each of the three directions. Table S.1 and Table S.2 in the supplementary quantify this by showing the three metrics (RMSE, Max abs. error, Standard deviation) over the interval [0, 0.3].

The same analysis is done for the *g_s_* parameter, where we see an improvement in the performance of Eucl-1-nn and Eucl-10-nn. Meanwhile, Apf-1-nn and Apf-10-nn, based on the APF-loss from Jæger et al. [22], have large fluctuations in estimation accuracy also for *g_s_*. Again, using the three metrics (RMSE, Max abs. error, Standard deviation), the difference in reliability is quantified; see Table S.1 and Table S.2 in the supplementary.

To further support these observations, Figure 7 shows the mean of the absolute error and the standard deviation for perturbation radii sampled uniformly on [0, 0.3] for each of the three directions A, B, and C. The first observation is that the error of StreaMRAK is significantly smaller than for the other methods both for the *g_Na_* and the *g_s_* estimations. More importantly, the small standard deviation exhibited by StreaMRAK, and to some degree, FALKON, illustrates the higher consistency of StreaMRAK compared to the other methods.

**Figure 7:**
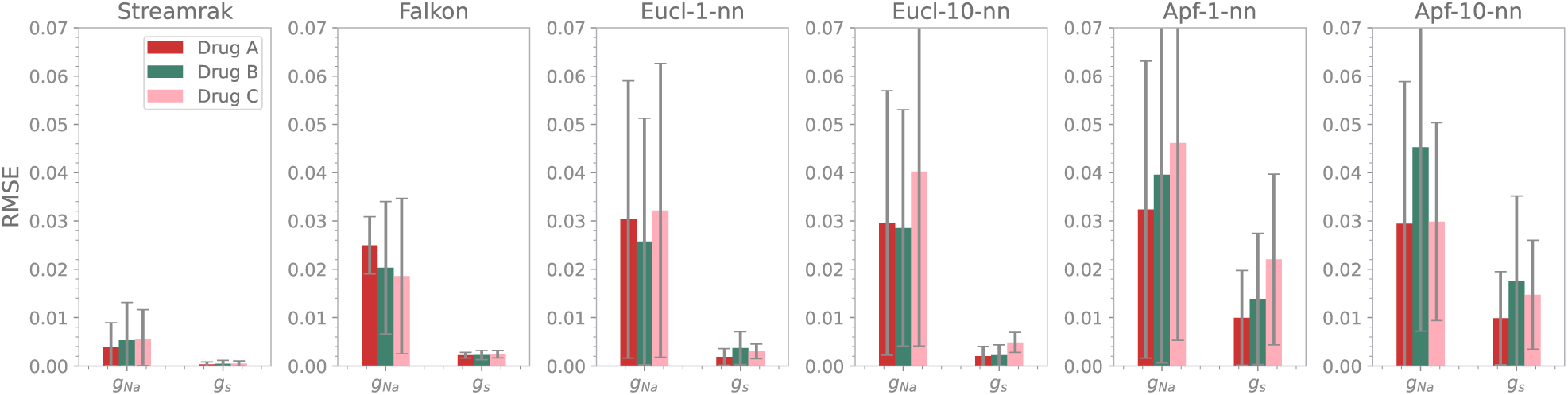
Figure showing the RMSE with error bars of the parameter estimations along three different perturbation directions on the interval [0, 0.3]. The perturbation directions correspond to the directions of Drugs A (red), B (green), and C (pink) defined in Figure 3.

Finally, for all methods, the error and variability are greater for the *g_Na_*parameter than for the *g_s_*parameter.

### 3.4 Experiment four: Demonstration of the inversion scheme as a tool for drug effect estimation using an example with simulated drugs

To demonstrate the use of StreaMRAK as a method for drug-effect estimation, we consider four simulated drugs [*A, B, C, D*] whose perturbation directions are indicated in Figure 3. Let *p_θ,r_* = (*θ, r*) be the polar representation of a parameter, centered at *p*_(0)_. For each drug, we run 100 experiments for a fixed angle *θ* and sample the perturbation radius *r* ∼ N(0.2, 1 × 10*^−^*^2^), namely a 20% perturbation with standard deviation ±5%.

All drugs considered in this experiment correspond to a perturbation from the unperturbed state *p*_0_ = (*g_Na_, g_s_*) = (1.0, 1.0), but in different directions. Drug A is a Ca^2+^-type current inhibitor (such as verapamil), drug C is a Na^+^ current inhibitor (such as quinidine), drug D is a Ca^2+^-type current agonist (such as BAYK8644), drug E is a Na^+^ current agonist (such as veratridine), and drug B is a mix of drugs A and C (such as a combination of quinidine and verapamil, or flecainide); [44, 45, 46, 47, 48, 49].

The estimation results for each of these drugs are summarised in Table 2 and Table 3. The root mean square error (RMSE) and the standard deviation (Std) are calculated over the 100 experiments. Meanwhile, the Pred column shows estimates of a specific perturbation whose magnitude is shown in the first row. Again, StreaMRAK outperforms the other methods with lower RMSE for all drugs. The estimation of the Na^+^ current parameter is consistently worse than the parameter for the slow inward current across all algorithms.

**Table 2:**
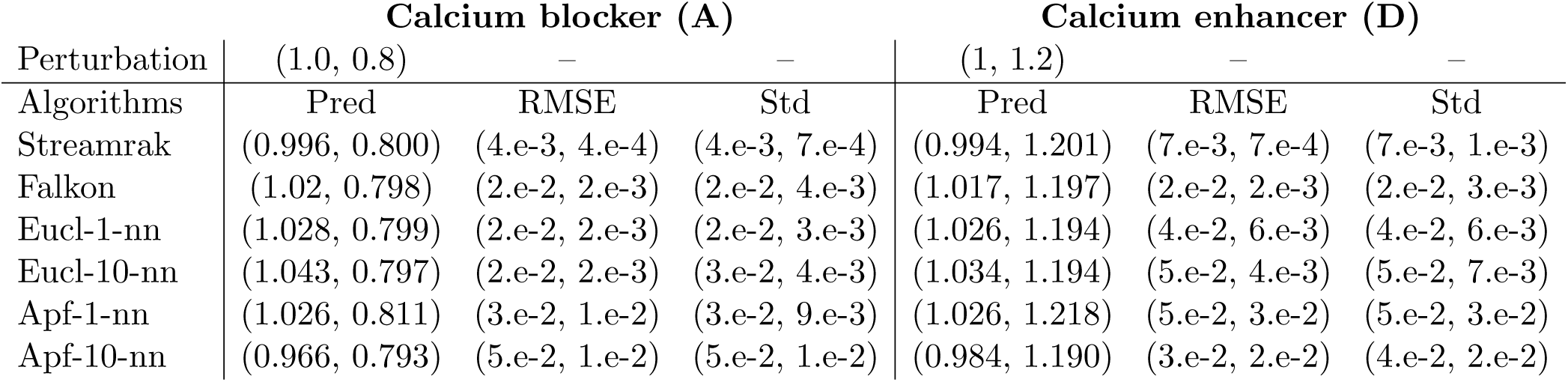
Estimation results of the algorithms w.r.t. the drug effects of the Ca^2+^ blocker, drug A, and the Ca^2+^ enhancer, drug D. The ”Pred” column is the predicted parameters (*p*_1_*, p*_2_), while ”RMSE” is the root mean square error of the estimation and ”Std” is the standard deviation.

**Table 3:**
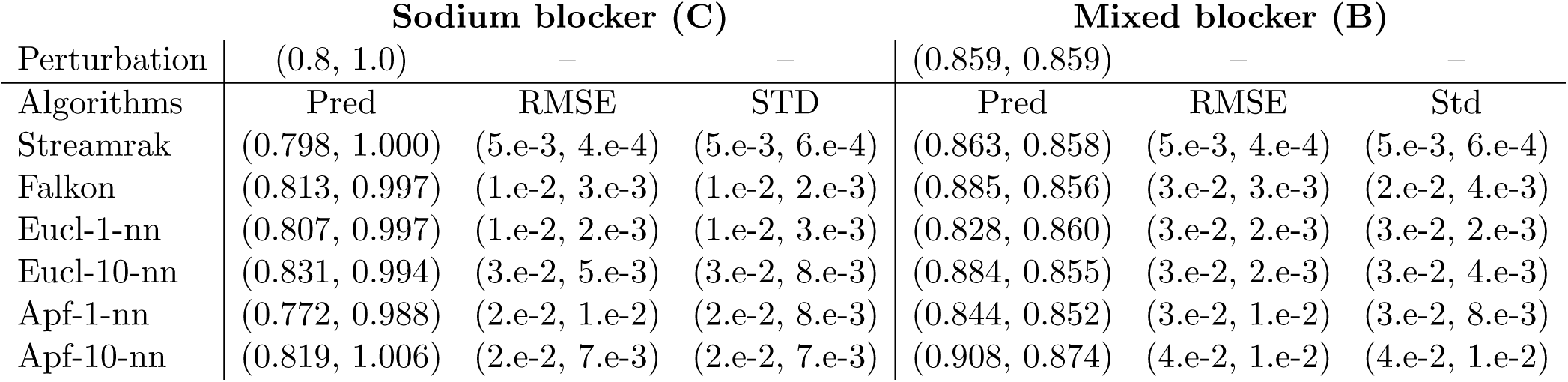
Estimation results of the algorithms w.r.t. the drug effects of the Na^+^ blocker, drug C, and the mixed current blocker, drug B. For column descriptions, see Table 2.

| Perturbation | Sodium blocker (C) |  |  | Mixed blocker (B) |  |  |
| --- | --- | --- | --- | --- | --- | --- |
|  | (0.8, 1.0) | – | – | (0.859, 0.859) | – | – |
| Algorithms | Pred | RMSE | STD | Pred | RMSE | Std |
| Streamrak | (0.798, 1.000) | (5.e-3, 4.e-4) | (5.e-3, 6.e-4) | (0.863, 0.858) | (5.e-3, 4.e-4) | (5.e-3, 6.e-4) |
| Falkon | (0.813, 0.997) | (1.e-2, 3.e-3) | (1.e-2, 2.e-3) | (0.885, 0.856) | (3.e-2, 3.e-3) | (2.e-2, 4.e-3) |
| Eucl-1-nn | (0.807, 0.997) | (1.e-2, 2.e-3) | (1.e-2, 3.e-3) | (0.828, 0.860) | (3.e-2, 2.e-3) | (3.e-2, 2.e-3) |
| Eucl-10-nn | (0.831, 0.994) | (3.e-2, 5.e-3) | (3.e-2, 8.e-3) | (0.884, 0.855) | (3.e-2, 2.e-3) | (3.e-2, 4.e-3) |
| Apf-1-nn | (0.772, 0.988) | (2.e-2, 1.e-2) | (2.e-2, 8.e-3) | (0.844, 0.852) | (3.e-2, 1.e-2) | (3.e-2, 8.e-3) |
| Apf-10-nn | (0.819, 1.006) | (2.e-2, 7.e-3) | (2.e-2, 7.e-3) | (0.908, 0.874) | (4.e-2, 1.e-2) | (4.e-2, 1.e-2) |

The Beeler-Reuter model is used to generate the AP traces corresponding to the predicted parameters. This provides an idea of which part of the AP gives rise to the estimation error. Figure 8 depicts how the AP traces relate to the trace of the actual perturbation (solid blue line). The AP trace corresponding to the unperturbed parameters is included for reference. There are, in particular, two regions where a discrepancy is present between the predicted and actual perturbations. These are the peak of the depolarization, mainly determined by the rapid Na^+^ current and the plateau before the final repolarization phase.

**Figure 8:**
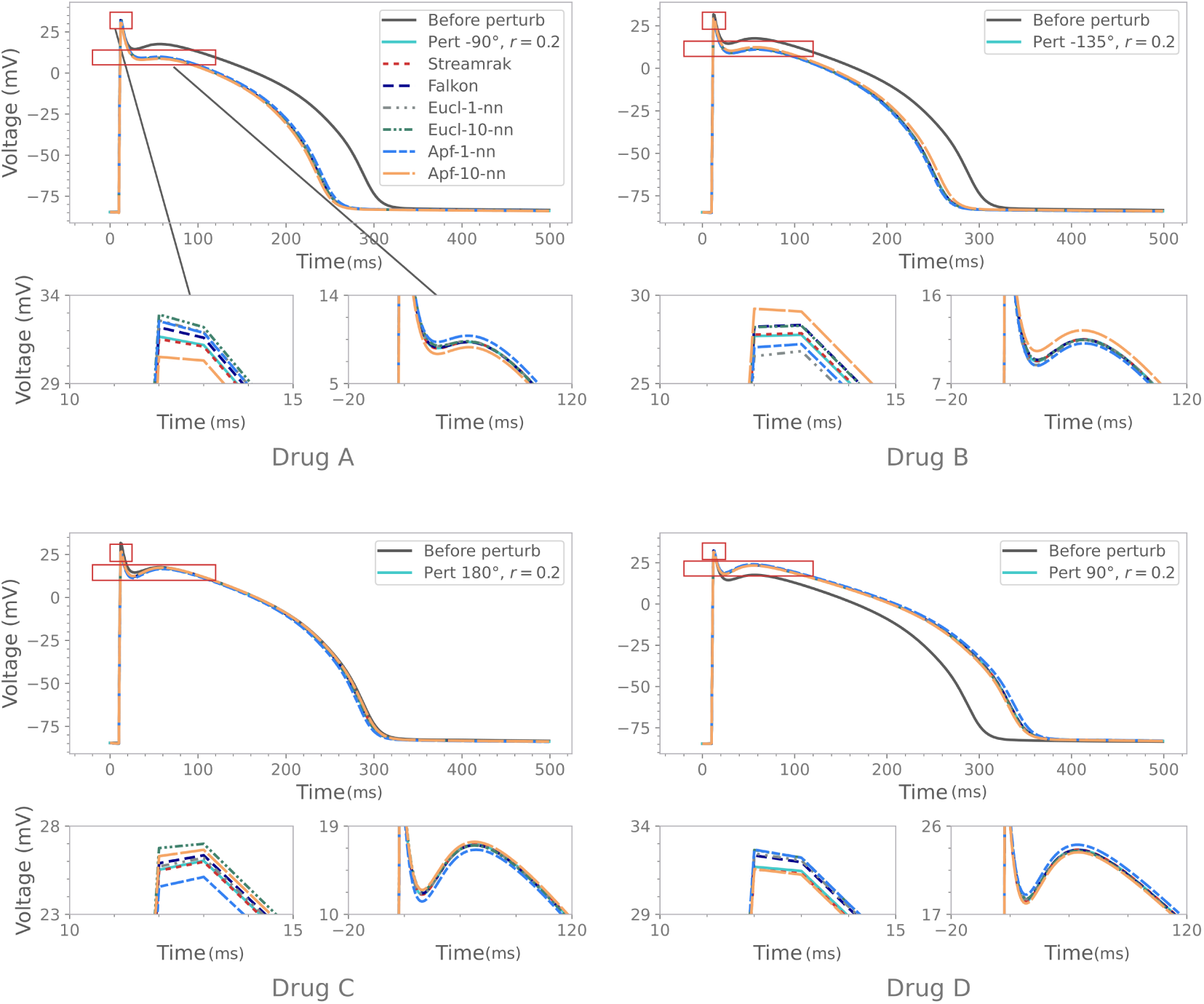
Comparison of how well the algorithms predict the drug effects of the drugs *A, B, C, D* from Figure 3. In each sub-figure, the upper panel shows the perturbation and the estimation from each algorithm. The unperturbed trace is included for reference. The lower panels highlight the first and the second ”peak” in the AP trace. The BR model is paced at 1 Hz with a sampling rate of 1 kHz. Only the first half of the pulse is included in the regression.

The Na^+^ inhibitor (Drug C) mainly affects the depolarization peak as seen from panel (Drug C) in Figure 8. Furthermore, the close-up in this panel shows that StreaMRAK gives a better fit to this peak compared to the other methods. Meanwhile, the Ca^2+^-type drugs A and D mainly affect the plateau and the repolarization phases, and these changes are more pronounced than the change in the depolarization peak. From the panels of drugs A and D, one can see that all methods give a close fit to the plateau and the repolarization phases. However, the close-up reveals that StreaMRAK has the tightest fit. Furthermore, the approximation of the depolarization peak is again better for StreaMRAK.

### 3.5 Performance analysis

To compare the overall performance and scalability of the methods, their parameter estimation capability and their time complexity are analyzed as a function of the training data size. The MSE is taken over all test estimations for each parameter to quantify the parameter estimation capability. The test data is the same as in Section 3.1 with 3000 points sampled uniformly from 𝒫 = [0.2, 2]^2^. Meanwhile, for the training data, a model is constructed on each of the sample sizes *n* ∈ [100, 500, 1000*, …,* 6000].

Consider the MSE over the test estimations for each parameter; the result is shown in Figure 9, where panel (A) is the MSE of the *g_Na_* estimations and panel (B) the MSE of the *g_s_* estimations. For both parameters, as the number of training samples increases, the MSE is reduced across all methods, and the error from StreaMRAK is consistently an order of magnitude lower than the rest. Furthermore, we see how StreaMRAK achieves a low estimation error with relatively small training sample sizes (2000 - 3000 samples). Meanwhile, the other methods do not reach a similar level of MSE even when doubling the sample size.

**Figure 9:**
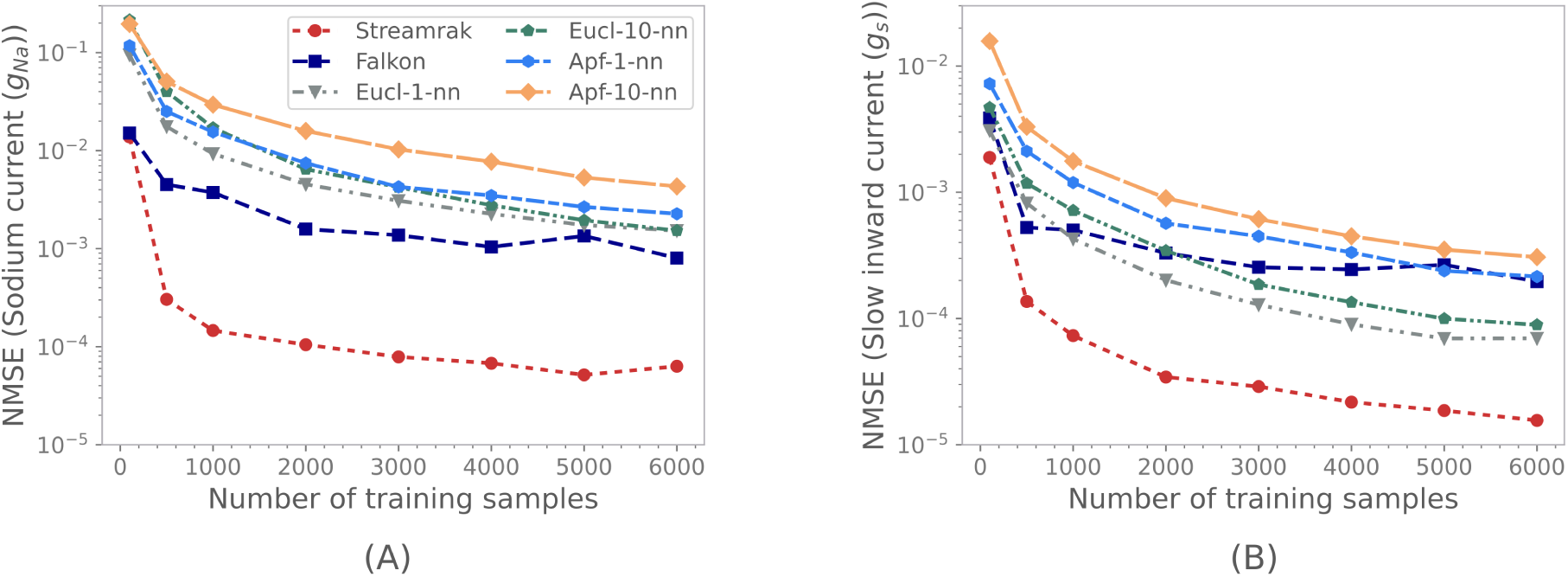
MSE of the estimations over the parameter domain *P* = [0.2, 2]^2^. Panel (A) shows the MSE of estimations of the *g_Na_* parameter (Na^+^ current). Panel (B) shows the MSE of estimations of the *g_s_* parameter (Ca^2+^-like current).

Figure 10 (A) compares the training time usage of the algorithms as a function of the training sample size. A notable advantage of the loss-minimization algorithms Eucl-1-nn, Eucl-10-nn, Apf-1-nn, and Apf-10-nn is that they do not require any training time because they only rely on ”memorizing” the training data. Although we need to consider the time required to construct the action potential features for each AP trace for Apf-1-nn, and Apf-10-nn, it is clear that the training time of StreaMRAK and FALKON is significant in comparison. However, as seen in Figure 9, StreaMRAK achieves a lower estimation error with far fewer samples. Because of this, the cost of generating the training samples must also be taken into account. This cost is indicated as the solid black line in Figure 10 (A), and it is clear that it is several magnitudes higher than the training times.

**Figure 10:**
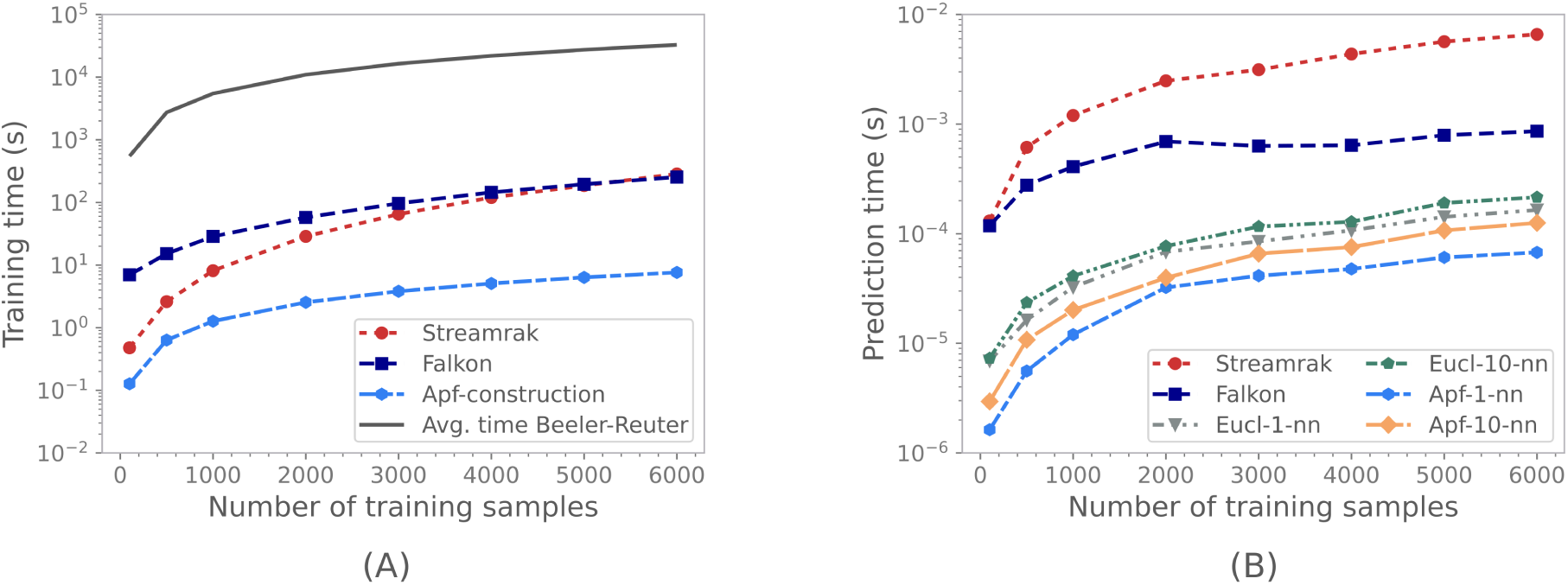
Comparison of time complexity of the algorithms. Panel (A) shows the training time of Streamrak and Falkon along with the time to construct the action potential features for the apf-k-nn algorithm. The solid dark line is the average time to solve for the AP traces of N samples using the Beeler-Reuter ODEs. Panel (B) shows the average estimation time, averaged over 2897 test samples.

Figure 10 (B), compares the average prediction (parameter estimation) time of each method as the model size (number of training samples) increases. Both StreaMRAK and FALKON are slower than the loss-minimization schemes.

## 4 Discussion

This study has compared two regression-based learning models, StreaMRAK and FALKON, with four nearest-neighbor loss-minimization schemes, Eucl-1-nn, Eucl-10-nn, Apf-1-nn, and Apf-10-nn. Our key findings are summarised in the following:

- Our experiments show that StreaMRAK estimates parameters with higher accuracy — roughly an order of magnitude higher — than both FALKON and the loss-minimization schemes.
- StreaMRAK, and to some degree FALKON, demonstrates a greater degree of reliability, both in terms of overall accuracy and consistency throughout the parameter domain.
- In particular, StreaMRAK demonstrates high reliability for the *g_Na_* parameter, which the loss-minimization schemes struggle with.
- Our experiments show that StreaMRAK requires significantly fewer training examples to reach a high estimation accuracy than both FALKON and the loss-minimization schemes.

In the following, we will discuss the results from Section 3 in more detail. In particular, we will focus on two key aspects: Reliability, which we discuss in Section 4.1, and computational performance, which we discuss in Section 4.2. We supplement this discussion with a note on the identifiability and sensitivity of currents in the Beeler-Reuter model; see Section 4.3. In Section 4.4 and 4.5, we make some special remarks with respect to the choice of model, the subset of parameters that we analyze, and the generalization of the inversion scheme. Finally, we consider related work, limitations, and future work in Sections 4.6 and 4.7.

### 4.1 Accuracy and reliability of methods

The experiments show that StreaMRAK performed the best regarding the absolute prediction accuracy and the consistency of the estimate on repeated tests. In particular, it has a higher estimation accuracy — roughly an order of magnitude higher — than both FALKON and the loss-minimization schemes Eucl-1-nn, Eucl-10-nn, Apf-1-nn, and Apf-10-nn. Furthermore, the parameter estimations of both StreaMRAK and FALKON are significantly more consistent across the parameter domain as demonstrated by Figure 4. These are the aspects of parameter estimation from cardiac AP inversion we sought to improve.

These findings show the advantage of using StreaMRAK as a tool for parameter estimation in the context of drug development. This is because, when exploring drug mechanisms, for example in heart-on-chip systems, the reliability of parameter estimates is vital for making consistent and accurate estimates of drug effects. In particular, an inversion scheme that provides high accuracy on average but has large variability over the parameter domain is undesirable because failure to estimate the effect of a particular drug dose can have severe consequences. An inversion scheme with high reliability is, therefore, a sensible requirement when using it for the purpose of drug-effect discovery. This study demonstrates that StreaMRAK satisfies this requirement.

Inspection of the estimation accuracy for different parameter combinations shows how the kernel methods StreaMRAK and FALKON are reliable when learning from training data where more than one parameter is responsible for the modulations of the AP traces and also more reliable with respect to estimating different parameters; See figure 5. This demonstrates that StreaMRAK and FALKON are better suited than the other loss-minimization schemes for scaling to larger training tasks from more complex models with several parameters.

Furthermore, Figure 6 and 7 demonstrate how StreaMRAK is reliable with respect to parameter estimation along specific directions in parameter space. This is an important property for an AP inversion scheme when the goal is to use it as a tool for drug calibration during a dose-escalation study, since measuring the response under a range of doses is an essential part of drug characterization. Increasing the dose of specific drugs results in perturbations in the parameter space with increasing magnitude away from the baseline biophysical cell. High reliability as a given parameter varies is, therefore, a crucial factor for adequate calibration. This study demonstrates that StreaMRAK satisfies this requirement.

As a final note, we mention that for estimating drug effects for clinical applications, it is necessary with more complex models and training sets with significantly more parameters (higher dimensional training data). What we have demonstrated in this study is that StreaMRAK has more potential for extension to larger models compared to the other inversion schemes considered in this study. In Section 4.5.1 we discuss in more detail the scalability of StreaMRAK to larger learning tasks from more complex models.

### 4.2 Computational considerations

Although the regression models StreaMRAK and FALKON have higher accuracy and are more consistent in their parameter estimation, these regression models come with an upfront training cost, as opposed to the loss-minimization schemes. Theoretically, this cost is 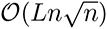 for StreaMRAK [31], where *L* is the number of resolution levels used in the model, and 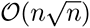 for FALKON [32]. For the APF-based methods, there is admittedly a cost associated with computing the action potential features for each AP trace. However, as seen in Panel (A) in Figure 10, this cost is significantly less than the training time of both StreaMRAK and FALKON.

Nevertheless, from panel (A) in Figure 9, we see that StreaMRAK, and to some degree, FALKON, requires significantly fewer training points to achieve a small MSE. In fact, from Panel (A) in Figure S.1 (See the Supplementary), we see the MSE as the number of training points is extended to *n* = 60000. Even then, for the *g_Na_* parameter estimation, StreaMRAK has almost an order of magnitude lower MSE. Meanwhile, for the *g_s_* parameter estimation, Eucl-1-nn and Eucl-10-nn catch up only after roughly *n* = 35000 training samples. This is somewhat expected because, with a high enough density of samples, any Voronoi cell (region closest to a point in a set than any other point in the set) will be approximately linear and small, which means minimizing the euclidean loss with the center of the Voronoi cell should have high accuracy. That StreaMRAK can do more for less is essential when we also consider the time it takes to construct the training samples by solving the AP model (Beeler-Reuter model). From Panel (A) in Figure 10, we see that solving the AP model has considerable time complexity. Furthermore, this time complexity is expected to increase for larger models. Consequently, the up-front training time is balanced by the reduced number of training samples for which the AP model must be solved.

Since the loss-minimization algorithms are based on minimizing a function over the training data, they must store all the training data, as the data is essentially their model. Therefore, the memory required for these algorithms is O(*n*). Furthermore, these models’ precision (resolution) is directly linked to the density of training samples. Consequently, because the number of samples necessary to cover a d-dimensional region grows exponentially with the dimension, the training set 𝒟*_n_* will need to be very large to make high-accuracy estimates for models with a large number of parameters.

The regression models, on the other hand, need the training data up-front to train the models. But once trained, significantly less memory is required for storage. For StreaMRAK this memory requirement is 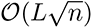, while for FALKON it is 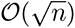. Smaller memory translates to easier transfer and implementation due to lower memory requirements for the host computer system. Furthermore, because these methods interpolate between the training samples, fewer training samples are required to achieve good estimates, as observed in panel (A) in Figure 9.

In terms of parameter estimation time (prediction time), we see from Panel (B) in Figure 10 that the regression-based methods are slower, which is consistent with analytical estimation time calculations. For StreaMRAK the analytical estimation time is 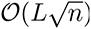, while for FALKON it is 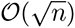. Meanwhile, using a KD-tree, the nearest-neighbor schemes have an estimation time of O(*d* log *n*) [50] (Using brute force, it is O(*dn*)). Here *d* is the intrinsic dimensionality of the AP traces (which is strongly related to the number of independent parameters in the model).

### 4.3 A note on the identifiability of currents in the Beeler-Reuter model

We use the identifiability analysis techniques discussed in Section 2.4 to gain a deeper understanding of the results from Section 3. In particular, we are interested in why estimates of the *g_s_* parameter have higher accuracy than the *g_Na_* parameter. Furthermore, we want to understand why the consistency of the *g_Na_* estimations is lower than for the *g_s_* parameter and why this is especially true for the loss-minimization methods.

The Beeler-Reuter model incorporates four currents of interest: the rapid sodium, slow-inward, time-dependent-outward current, and time-independent outward current, whereof the first two are parameterized by respectively the *g_Na_* and *g_s_* parameters in this study. Using the spectral analysis from Jæger et al. [27], as described in Section 2.4.1, we can analyze the sensitivity and identifiability of these currents in the Beeler-Reuter model. Let *e* = (1, 0, 0, 0) represent the sodium current, *e* = (0, 1, 0) the slow inward current, *e* = (0, 0, 1, 0) the time-dependent outward current, and *e* = (0, 0, 0, 1) the time-independent outward current. From the analysis, we find the singular values *σ* = (*σ*_1_*, σ*_2_*, σ*_3_*, σ*_4_) = (707.8, 6.9, 0.18, 0.066) with corresponding singular vectors

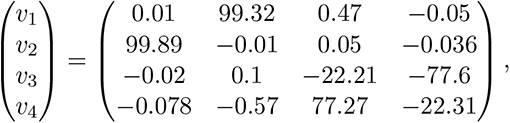

where the singular vectors are multiplied by a factor of 10^2^ to enhance readability.

This means the current parameterized by *g_s_* projects mainly along the largest eigenvalue direction, while *g_Na_* is projected mainly along *v*_2_ with singular value *σ*_2_ ≪ *σ*_1_. Meanwhile, the time-dependent and time-independent outward currents both project along *v*_3_ and *v*_4_. Jæger et al. [27] observed that the change in the AP traces when perturbing along a singular vector is proportional to the corresponding singular value. Consequently, this analysis shows that for the Beeler-Reuter model, we can expect less sensitivity for the *g_Na_* parameter than for the *g_s_* parameter, which helps explain why the accuracy in estimating *g_Na_* is lower than for *g_s_*.

To supplement this analysis, we consider the geometrical analysis proposed in Section 2.4.2. We let the parameter domain Ω be the ball B(*p*_0_*, δ*) ∈ 𝒫 centered at *p*_0_ = (1, 1) with radius *δ* = 0.2. Panel (D) in Figure 11 illustrates this ball. The corresponding AP traces lies on a 2-dimensional surface in 𝒱*_T_* ∈ R*^T^*, which we embed in 3 dimensions using Laplacian eigenmaps as shown by the yellow surface in Panel (A) in Figure 11. The grey ellipse in panel (A) indicates the embedded AP traces corresponding to the parameters on the blue circle in panel (D).

**Figure 11:**
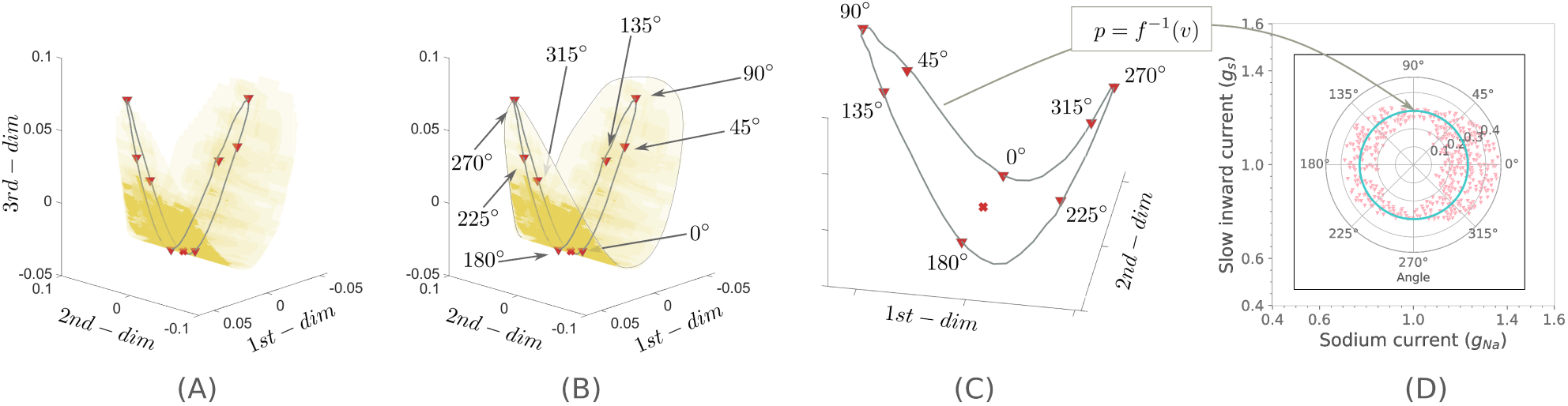
Panel (A) shows the 3-dimensional projected embedding of a ball in the space of action potential curves *V* (Embedding using the Laplacian Eigenmaps algorithm [40]). The grey ellipse in (A) corresponds to a circle of radius *r* = 0.1 centered at *p* = (1, 1) in the parameter space. Panel (B) indicates how the directions in the parameter space map to the embedding. Panel (C) and (D) illustrate the relationship between a perturbation ring of radius *r* = 0.3 in parameter space (light blue circle in panel(D)) and the corresponding ellipse in *V* (grey ellipse in panel (C)). The pink triangles correspond to voltage curve samples from a band around the ellipse in panel (C) that are mapped back into the parameter space.

From Section 6.4 in the appendix, it is clear that the shape of the ellipse directly relates to the derivative of the inverse map *f^−^*^1^. Along the *g_Na_* axis, the ellipse is narrow, and therefore the derivative is large. Meanwhile, along the *g_s_* axis, the derivative of *f^−^*^1^ is comparatively small. Clearly, high accuracy in voltage space is particularly vital in perturbation directions where small changes in AP-trace space significantly affect the underlying parameters. To illustrate this, consider AP traces from a narrow band along the ellipse. The corresponding parameters are shown as pink triangles in Panel (D). What is apparent is that, along the *g_Na_* axis, the spread in parameters is considerable compared with the spread along the *g_s_* axis. Consequently, because minor errors when matching the AP traces lead to substantial changes in the estimates of *g_Na_*, we can expect the variability in the *g_Na_* estimations to be more significant than for the *g_s_* estimations.

The difficulty of learning the effect of the *g_Na_* parameter is illustrated in Figure 12, which shows the AP traces as they vary when the parameters are perturbed along different directions from the initial *p*_0_ = (1, 1). Clearly, the *g_Na_* parameter only leads to subtle changes in the AP traces, primarily affecting the depolarization peak. Consequently, if a learning model does not capture these subtle differences, it will lead to reduced estimation accuracy, which is precisely what we observe for the *g_Na_* parameter.

**Figure 12:**
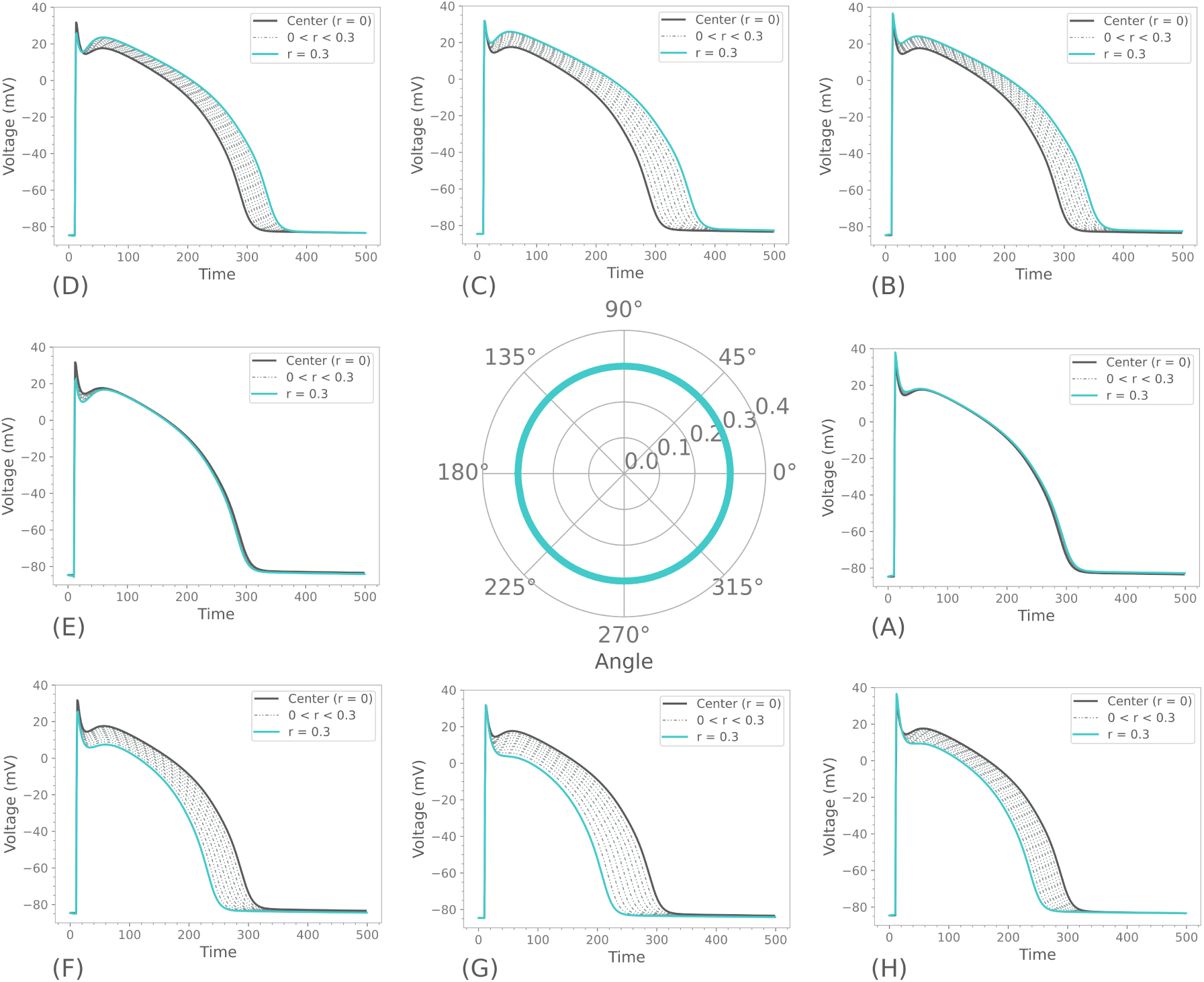
Illustration of how the voltage curves change as a function of direction and radius away from the center *p* = (1, 1). The middle panel is a polar representation of the direction and radial distance away from *p* = (1, 1). Each panel (A)-(H) shows 10 curves equally spaced on the interval [0, 0.3]. In particular, the center curve *r* = 0 is a dark solid line, and the curve at *r* = 0.3 is a solid light-blue line. Note that panel (A) corresponds to direction 0*^◦^*, (*B*) to direction 45*^◦^* etc. The BR model is paced at 1 Hz with a sampling rate of 1 kHz. Only the first half of the pulse is included in the regression.

For example, we consider the drug estimation in Figure 8, where we see that StreaMRAK is better at detecting critical features of the AP traces. For instance, consider the following example. A key identifier of Na^+^ current inhibitors (such as quinidine) is the peak of depolarization. This drug effect can be simulated by perturbations along both C and B in Figure 3, and we see from Figure 8 that in both cases, StreaMRAK succeeds in capturing the depolarization peak, while the other methods fail to do so. Consequently, a significant error is observed in these methods’ *g_Na_* estimation by failing to distinguish differences in this peak.

### 4.4 A note on the choice of parameters and model for the study

A limitation of using the 1977 Beeler-Reuter model is that it does not incorporate the presence of more specific inward and outward currents that are present in contemporary models and would introduce challenges with identifiability due to overlapping effects. These are challenges that warrant further investigation in order to extend our proposed methods to more clinically relevant scenarios; However, the focus of this study has been to compare StreaMRAK to existing schemes with respect to reliability across the parameter domain. Meanwhile, the analysis of identifiability issues that arise from overlapping current effects is left for a separate study. Because of this, the use of the 1977 Beeler-Reuter model is a natural choice for the purpose of this study.

This study did not include parameter estimations for the time-independent potassium K^+^ current and the time-dependent outward current. These parameters are interesting from a drug screening perspective and carry important clinical implications. However, for the purpose of this study, we have focused on rigorous analysis of a subset of parameters which enabled an interpretable and thorough comparison of different inversion schemes across the parameter domain. For demonstrating StreaMRAK as a tool for estimating drug effects with clinical utility, it will be necessary to consider more elaborate AP models with significantly more parameters, complexity, and challenges, such as Tusscher and Panfilov [9] and O’Hara et al. [11].

### 4.5 Generalization of the inversion scheme

We discuss the scalability of the inversion scheme to larger models and the generalization to more complex AP models. We also offer our thoughts on how the method will perform on AP traces recorded at multiple pacing rates and different time resolutions.

#### 4.5.1 Scalability to larger models

The scalability of the kernel methods StreaMRAK and FALKON, to larger data sets with higher dimensionality (more parameters) has already been demonstrated in several studies [32, 51, 31]. Furthermore, because the learning algorithms are non-parametric, they do not rely on specific information about the AP trace and AP model at hand. As such, they should also work on training data generated by more complex AP models.

We note that as the number of parameters in the training data increase, one needs more training samples to maintain the estimation accuracy. This problem is known as the *curse of dimensionality* [52], and is a well-known reality of learning that also applies to the loss-minimization schemes Eucl-1-nn, Eucl-10-nn, Apf-1-nn, and Apf-10-nn. What we have shown in this study is that StreaMRAK requires significantly fewer training samples to achieve high accuracy for a given number of parameters. Therefore, StreaMRAK will be the better option when considering more complex models with more parameters.

#### 4.5.2 Differences in AP recordings

For AP traces recorded with different recording systems, we distinguish between two cases: The first is AP traces with different pacing rates, which is relevant when studying, for example, anti- and proarrhythmic drug effects. The second is AP traces recorded at different sampling frequencies, which is often the case for different experimental setups and recording systems. For example, microelectrode arrays typically sample at frequencies in the order of 10 − 20 kHz [5, 53], while live cell fluorescence microscopy sample at significantly lower frequencies *<* 1kHz.

Regarding the first case. The pacing rate can affect channel kinetics and, therefore, the relationship between the AP trace and the underlying parameters. This means the inverse map *f^−^*^1^ we are approximating will differ depending on the pacing rate. As such, a model trained on a set of AP traces with the same pacing rate (as done in this study) is only representative of current parameters for AP traces with this pacing rate (or very similar pacing rates). We note that this would also be the case for any other learning scheme since changing the pacing rate alters the target function to learn. Because of the importance of pacing rate with respect to detecting antiarrhythmic effects, future studies should consider the performance of StreaMRAK in this regard. The extension of StreaMRAK to parameter estimation from multiple pacing rates can be done straightforwardly. The first step is to include more than one pulse of the AP trace; In Cairns et al. [20], this was done by including two pulses, which should be sufficient. To allow the traces to be compared, the two-pulse AP traces must be embedded in the same vector space, as explained in Section 2.2.4. The next step is to train the model on these two-pulse AP traces using a selected set of relevant pacing rates S. The performance of the inversion scheme should then be explored with respect to parameter estimations of AP traces with pacing rates both in S and outside of S.

For clinical applications involving *in-vitro* measurements of AP traces, the relevant pacing rates are determined by the practitioner in the lab, which measures the AP traces at pacing rates relevant to the problem that is studied. To invert these traces, the model should be trained at the same set of pacing rates. For purely *in-silico* experiments, the pacing rates used for training the model should, in a similar manner, be relevant to the problem that is studied.

Regarding the second case. As noted in Section 2.2.4, a requirement for the proposed method is that the sampling frequency of the training data and the AP traces measured *in-vitro* must be the same. Otherwise, the AP trace would not be comparable. However, should the time grids differ due to different sampling frequencies, it is possible to align them by time interpolation or downsampling, which is possible because the AP traces are smooth functions of time. This way, one can avoid training the model again due to differences in sampling frequency. Furthermore, in this study, we have demonstrated the method on synthetic AP traces with a sampling rate of 1 kHz. However, nothing prevents the method from being trained on any other reasonable sampling rate.

### 4.6 Complementary studies

This study has demonstrated StreaMRAK as a computationally efficient AP trace inversion scheme with greater reliability than alternative schemes that have been previously applied for this purpose [30]. As such, our contribution is a piece in a larger effort to develop efficient drug-effect-estimation pipelines based on high-throughput, optical measurements of AP traces.

Tveito et al. [30] demonstrate, using synthetic and experimental data, how optically measured AP traces from human induced pluripotent stem cell-derived cardiomyocytes (hiPSC-CMs) can be inverted to gain information on drug effects in mature ventricular cells. This scheme has implications for decreasing dependency on highly specialized laboratory expertise, which can contribute to reducing the cost and time required to develop and launch a new drug to market. The relevance of our method can be seen in relation to the utility of this pipeline.

Other recent studies have also contributed to advancing the inversion step. In particular, Jæger et al. [22] introduces a new computational scheme for inverting the AP model based on a continuation-based optimization algorithm [23], which searches for the optimal parameter of an AP trace by moving iteratively from a known initial parameter. Meanwhile, Jæger et al. [54] show in the context of hiPSC-CMs that by combining the extracellular potential with the membrane potential and the calcium concentration, the identifiability of the fast sodium (Na^+^) current, the main contributor to the rapid upstroke of the AP, can be improved. They also show that other significant currents can be identified in this way.

Several alternative pipelines using hiPSC-CMs measurements to predict drug-effect responses in adult human ventricular cardiomyocytes have also been developed. Using statistical modeling and multivariable regression, Gong and Sobie [55] developed a cross-cell type regression model for predicting responses in human adult ventricular cardiomyocytes from measurements in hiPSC-CMs. Meanwhile, Paci et al. [56] combined *in silico* simulation trials with optical measurments of AP and Ca^2+^ traces in hiPSC-CMs to estimate drug effects. Similarly, Passini et al. [57] estimated drug effects by combining *in silico* simulation trials with Ca^2+^ trace measurements from hiPSC-CMs and also from electrocardiogram measurements from animal preparations.

The use of machine learning techniques for drug effect estimation has also been used in several studies. Lancaster and Sobie [58] developed a classifier for estimating drug torsadogenicity using statistical analysis and machine learning techniques. Extracting several key features from the AP and intercellular Ca^2+^ traces of clinical data, a support vector classifier was trained to distinguish between arrhythmogenic and non-arrhythmogenic drugs. Aghasafari et al. [59] developed a deep-learning classification network capable of learning from time-series data, such as cardiac AP traces. The network was used to differentiate between drugged and drug-free adult ventricular myocytes from measurements of AP traces in hiPSC-CMs and to estimate how electrophysiological perturbations influence the maturation of hiPSC-CMs. The network resembles our proposed scheme as it requires no assumptions about system parameters and can, in principle, be used for any time series data.

### 4.7 Limitations and future work

We have demonstrated the parameter estimation capability of StreaMRAK on the 1977 Beeler-Reuter AP model of mammalian ventricular cardiomyocytes [34]. Although the BR model captures currents that are of interest for studying the electrophysiological effects of drugs, such as the fast inward sodium (Na^+^) current and a slow inward current, primarily carried by calcium (Ca^2+^) ions, there exist several other AP models which incorporate other more specific currents. Notable examples include: the Ten Tusscher ventricular model, which incorporates several other transmembrane currents, transmural differences in currents, and more complex calcium dynamics [9]; the Grandi model which includes an excitation-contraction component [10]; and the O’Hara (ORd) model which more accurately models the human-specific undiseased ventricular action potential and causes of arrhythmic behavior [11].

These models capture more detailed electrophysiology and give a better representation of the cardiac cell, albeit less interpretable. Because of this, future work should focus on extending the analysis in this paper to these models. Of particular importance with these more complex models is the presence of more realistic potassium (*K*^+^) current dynamics, which is a key current when assessing anti- and pro-arrhythmic drugs [60]. Furthermore, with more complex models come more involved identifiability problems; sensitivity problems due to subtle (hard to detect) changes to the AP, such as observed in this study with the *g_Na_* parameter; And identifiability problems due to parameters with overlapping effects on the AP that either significantly reduce or cancel each other out [26].

For the extension to more complex AP models, it will be necessary with an in-depth study of the performance of StreaMRAK in the context of the second of these identifiability problems. In particular, one should aim at isolating a subset of parameters from a more complex model for which this problem occurs, similar to what has been done in this study with respect to scalability, reliability, and *g_Na_*.

A notable drawback with the proposed kernel regression schemes of StreaMRAK and FALKON is their considerable upfront training time compared to the loss-minimization schemes that do not require any training in advance. However, as shown in this study, this is balanced by the reduction in sample points required to achieve high accuracy. This is because reducing the number of training points reduces the computational burden of solving the system of ODEs in the AP model and the memory requirement of storing the samples. This reduction is significant due to the expense of solving the ODE compared to the training time, as seen from Figure 10. Nevertheless, it is worth mentioning that in situations where an extensive upfront training time is undesirable, and high reliability and low memory are of less importance, then the loss-minimization schemes can be more suitable options.

Furthermore, our experiments demonstrate StreaMRAK on an AP model for adult human cardiomyocytes. Future work should aim to demonstrate this model in the drug-effect-identification pipeline developed in Tveito et al. [30]; Jæger et al. [22] and to extend the demonstration to AP models of immature human induced pluripotent stem cells (hiPSC-CMs), such as Paci et al. [36, 61]. The generalization to AP models of hiPSC-CMs is straightforward, as StreaMRAK is a non-parametric learning scheme that does not rely on model-specific input about the training data.

Moreover, validation of the methodology against experimental drug response data should be performed. Initially, in terms of drug response experiments on hiPSC-CMs in microphysiological systems, since experimental control data for hiPSC-CMs are more readily attainable. However, in future work, we also aim to experimentally validate the StreaMRAK inversion for AP models of adult human cardiomyocytes. Drug response data for adult human cardiomyocytes is difficult to attain because both low-throughput patch clamp techniques and higher-throughput, multi-cell platforms require highly specialized laboratory expertise and high initial expense for the instrumentation [1, 2]. Because of this, validation of the inversion of AP models of adult human cardiomyocytes will mainly be done using live cell fluorescence microscopy [3, 4]. This can be done by feeding the inverted parameters back into the AP model and then comparing the resulting AP traces with the measured ones.

## 5 Conclusion

In this study, we have introduced StreaMRAK as a new tool for inverting cardiac AP traces using a subset of parameters from the 1977 Beeler-Reuter model. We have demonstrated that StreaMRAK is a scalable method that offers greater reliability than existing inversion schemes, especially for AP model parameters with effects that are hard to detect. This systematic study offers a foundation for applying StreaMRAK to larger and more complex AP models with clinical value.

## Acknowledgments

AO is funded by Simula Research Laboratory. AC was partially funded by NSF DMS 1819222 and 2012266, and a gift from Intel research. NF is funded by Simula Research Laboratory.

## 6 Appendix

We recapture some minor results and definitions and finish with a closer look at kernel methods and a more extensive outline of StreaMRAK and FALKON.

### 6.1 The inverse of multivariate vector functions

The inverse function theorem 6.1 states that a vector-valued function over multiple variables *f* : *X* ⊆ ℝ*^N^* → ℝ*^N^* has a local inverse at **x**_0_ ∈ *X* if the jacobian *J_f_* (**x**_0_) is invertible. In other words, if the determinant det *J_f_* (**x**_0_) ̸= 0.

**Theorem 6.1** (The inverse function theorem). (See for instance Krantz et al. [62]) Let X ∈ R*^n^ be open, and let f* : *X* → R*^n^ be a continuously differentiable function f* ∈ *C*^1^(*X,* ℝ*^N^*). Let **x**_0_ ∈ *X and let J*(**x**_0_) be the Jacobian at **x**_0_*. If J_f_* (**x**_0_) *is invertible (i.e.* det *J_f_* (**x**_0_) ̸= 0*), then there exists an open neighborhood* N(**x**_0_) *such that the inverse f^−^*^1^ *of f* : N(**x**_0_) → *f* (N(**x**_0_)) *exists and J_f −_*_1_ (*f* (**x**_0_)) = (*J_f_* (**x**_0_)).

### 6.2 Definitions and minor results

**Definition 6.2** (Jacobian). Consider a function f : *X* ⊆ ℝ*^N^* → R^m^ where all partial derivatives exists for **x** ∈ *X. Then the Jacobian matrix of f is defined as*

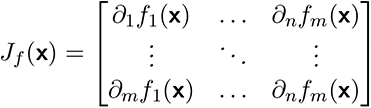

**Definition 6.3** (Positive semi-definite matrix). Let **A** ⊂ R^n×n^ be a symmetric matrix. If **x**^⊤^**Ax** ≥ 0 for all **x** ∈ R^n^ then we say that **A** is a positive semi-definite matrix.

**Definition 6.4** (Positive definite function). Let f : R → C be a complex-valued function and **A**_ij_ = *f* (**x**_i_ − **x**_j_) be the symmetric matrix induced by f . If **A** is a positive semi-definite matrix as follows from Def. 6.3, then we say f is a positive definite function.

**Definition 6.5** (Epsilon cover). Consider the metric space (X *, d*) *and let* Γ*_ε_*⊂ X *be a subset of samples from* X *. Furthermore, let ϵ, δ >* 0 be two constants. If for every sample **x**^′^ ∈ Γ*_ε_ we have ε < d*(**x**^′^, **x**) *< ε* + δ for all **x** ∈ Γ*_ε_ such that* **x** ≠ **x**^′^. Then we call Γ*_ε_ an epsilon cover of* X.

### 6.3 A note on invertability

From the inverse function theorem 6.1, we know that a multi-variable function such as *F_T_* is locally invertible over a region where the jacobian *J_F_* (*p*) from Def 6.2 (the matrix containing all partial derivatives of *F_T_*) is invertible, i.e., det *J_F_* (*p*) ≠ 0. This means, in essence, that we can construct a model of *F^−^*^1^ from some domain V to the parameter space 𝒫 provided the forward map is unique; all voltage curves *v* ∈ 𝒱*_T_* correspond to unique parameters *p* ∈ P. We note that if a domain 𝒱*_T_* contains voltage curves *v_i_* ≠ *v_j_* for which the corresponding parameters are equal *p_i_* = *p_j_*, it is necessary first to split the domain into sub-domains over which the forward map is unique. Separate models can then be learned for each of these sub-domains.

### 6.4 Local distances in the image of a non-linear map

Consider the spaces *X, Y* and let *f* : *X* → *Y* be a non-linear map between them. Let *x*_0_ ∈ *X* be a point in *X* and *y*_0_ = *f* (*x*_0_). We then have that the ball *B_X_*(*x*_0_*, δ*) in *X*, defined as

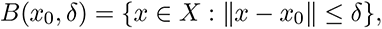

maps to the ellipse

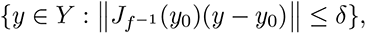

in the output space *Y*, where *J_f −_*_1_ (*y*_0_) is the Jacobian (derivative) of *f^−^*^1^ at *y*_0_, see section 6.1 in Singer and Coifman [63] for a proof.

### 6.5 Kernel methods

The use of kernel methods for supervised learning has a strong theoretical foundation [64, 65, 66]. Kernel methods rely on the use of a positive definite kernel *k* : *X* ×*X* → *R*, with X ⊂ R*^d^*, to construct an infinite-dimensional vector space of functions 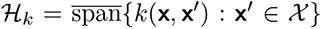, called a reproducing kernel Hilbert space (RKHS). See Def 6.4 for the definition of a positive definite function. As *H_k_* is a vector space, any function *h* ∈ *H_k_* can be expressed as a linear combination of the basis vectors *k***_x_**(**x***^′^*) := *k*(**x**, **x***^′^*). Furthermore, the advantage of creating this function space comes to light through the ”kernel trick” [67, 68], which states that each basis vector corresponds to the inner product between two non-linear feature vectors *ϕ*(**x**)*, ϕ*(**x***^′^*) ∈ H*_k_*, namely *k***_x_***′* (**x**) = ⟨*ϕ*(**x**)*, ϕ*(**x***^′^*)⟩. As the feature vectors can be highly non-linear functions in **x**, a linear model in H*_k_* corresponds to a highly complex function in the input space X, which explains the great expressive power of this space.

Consider training data 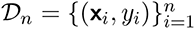 drawn from the sample space *X* × *R* according to some probability distribution *ρ*. Supervised learning aims to train a model that gives a good approximation of the function that generated this data. Using an RHKS as our model space, we can then formulate the optimization problem

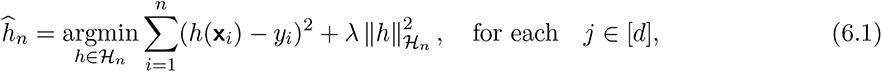

where the model space is restricted to the finite-dimensional subspace H*_n_* ⊂ H*_k_* due to the finite size of the training data. The solution to this problem is guaranteed by the Representer theorem [69] to be a linear combination of the basis vectors that make up H*_n_*, namely

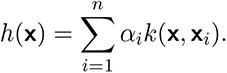

It follows that the optimization problem in Eq. (6.1) can be solved as the linear system (**K**+*λ***I***_n_*)***α*** = **y**, where **K***_ij_* = *k*(**x***_i_,* **x***_j_*), *y* = (*y*_1_*, …, y_n_*)*^⊤^* and ***α*** = (*α*_1_*, …, α_n_*)*^⊤^*. However, inverting the *n* × *n* matrix **K** is computationally expensive, with time complexity of O(*n*^2^). To remedy this, the large-scale kernel method FALKON [32] introduces several improvements to speed up the inversion.

#### 6.5.1 FALKON

In FALKON the computation complexity of KRR is reduced by selecting a random subset of the training data 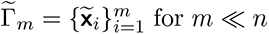 for *m* ≪ *n*, which we refer to as landmarks or Nyström sub-samples. The model space is then reduced to the span of the kernels centered on these sub-samples, namely 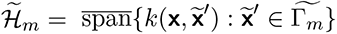. Solving the optimization problem in Eq. (6.1) with the reduced model space gives rise to a linear system involving the much smaller *n* × *m* kernel matrix **K***_ij_* = *k*(**x***_i_,* **x***_j_*). Further improvements are made by introducing a pre-conditioner and solving the linear system iteratively using conjugate gradient with early stopping. For more details on the algorithm, we refer to Rudi et al. [32] and Oslandsbotn et al. [31].

#### 6.5.2 StreaMRAK

The streaming multi-resolution adaptive kernel algorithm StreaMRAK introduced in Oslandsbotn et al. [31], significantly improves the model used in FALKON and standard KRR. Using principles from boosted gradient descent, an iterative model is introduced which solves the KRR over several levels. The iterative model is on the form

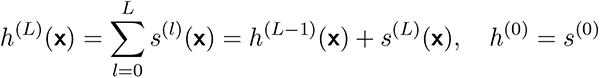

where

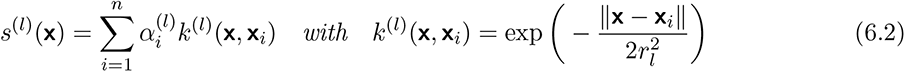

is a correction term to the previous level. The coefficients *α_i_*^(^*^l^*^)^ at each level *l* are found by using the FALKON solver to regress on the residuals **d**^(^*^l^*^)^ = (*d_i_*^(^*^l^*^)^*, …, d_n_*^(^*^l^*^)^) ∈ *R^n^* from the previous level. The residuals are defined as

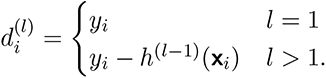

where *h*^(*l*^*^−^*^1)^(**x***_i_*) is the prediction of *y_i_*using the trained model from the previous level.

In StreaMRAK the kernel bandwidth *r_l_* is reduced at each new level as *r_l_* = 2*^−l^r*_0_. Furthermore, instead of the random sub-sampling of landmarks used in FALKON, the landmarks are selected from an epsilon cover with epsilon *ε* = *r_l_*, see Def. 6.5. This way the distance between landmarks is tailored to the kernel bandwidth, which allows the optimal utility of each sub-sample. With these two choices, StreaMRAK implements an adaptive multi-resolution scheme that is more robust at learning complex functions than standard KRR solvers and requires significantly less memory [31].

## Supplementary

We include two experiments that did not fit in the main text. In Figure 13, we see the MSE of the predictions over the parameter domain *P* = [0.2, 2]^2^ as a function of training data size *n*, where *n* ∈ [1000, 60000]. In Table 4 and Table 5 we see the Root mean square error (RMSE), Max absolute error (Max abs. err.) and standard deviation (Std) for the parameter estimations of (*g_Na_, g_s_*) along the drug directions A, B and C from the experiment presented in Section 3.4.

**Figure 13:**
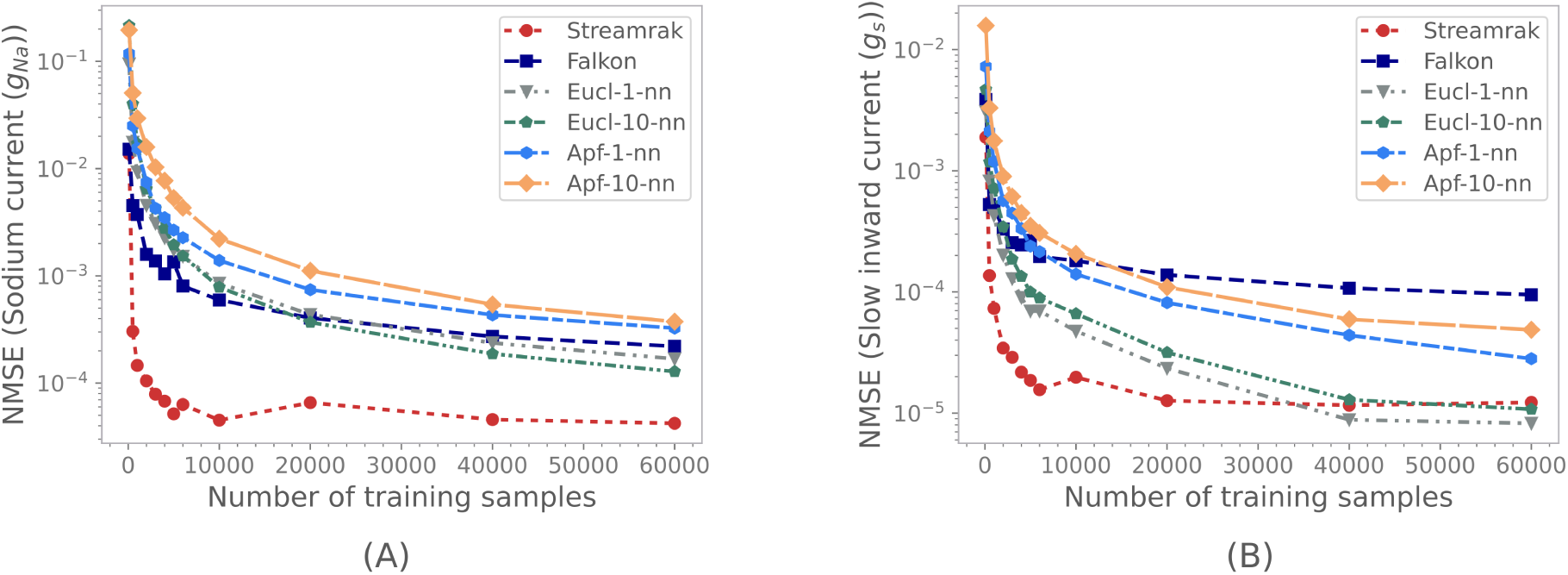
MSE of the predictions over the parameter domain *P* = [0.2, 2]^2^ as a function of training data size. Panel (A) shows the MSE of predictions of the *g_Na_* parameter (Na^+^ current). Panel (B) shows the MSE of predictions of the *g_s_* parameter (Ca^2+^ -like current).

**Table 4:**
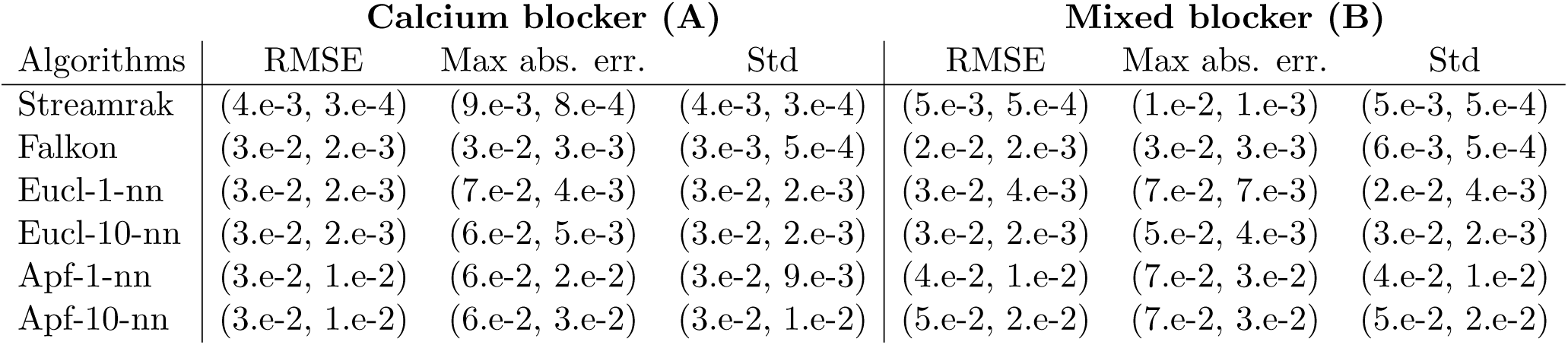
Table showing Root mean square error (RMSE), Max absolute error (Max abs. err.) and standard deviation (Std) for the parameter estimations of (*g_Na_, g_s_*) along the drug directions A and B from the experiment presented in Section 3.4.

| Algorithms | Calcium blocker (A) |  |  | Mixed blocker (B) |  |  |
| --- | --- | --- | --- | --- | --- | --- |
|  | RMSE | Max abs. err. | Std | RMSE | Max abs. err. | Std |
| Streamrak | (4.e-3, 3.e-4) | (9.e-3, 8.e-4) | (4.e-3, 3.e-4) | (5.e-3, 5.e-4) | (1.e-2, 1.e-3) | (5.e-3, 5.e-4) |
| Falkon | (3.e-2, 2.e-3) | (3.e-2, 3.e-3) | (3.e-3, 5.e-4) | (2.e-2, 2.e-3) | (3.e-2, 3.e-3) | (6.e-3, 5.e-4) |
| Eucl-1-nn | (3.e-2, 2.e-3) | (7.e-2, 4.e-3) | (3.e-2, 2.e-3) | (3.e-2, 4.e-3) | (7.e-2, 7.e-3) | (2.e-2, 4.e-3) |
| Eucl-10-nn | (3.e-2, 2.e-3) | (6.e-2, 5.e-3) | (3.e-2, 2.e-3) | (3.e-2, 2.e-3) | (5.e-2, 4.e-3) | (3.e-2, 2.e-3) |
| Apf-1-nn | (3.e-2, 1.e-2) | (6.e-2, 2.e-2) | (3.e-2, 9.e-3) | (4.e-2, 1.e-2) | (7.e-2, 3.e-2) | (4.e-2, 1.e-2) |
| Apf-10-nn | (3.e-2, 1.e-2) | (6.e-2, 3.e-2) | (3.e-2, 1.e-2) | (5.e-2, 2.e-2) | (7.e-2, 3.e-2) | (5.e-2, 2.e-2) |

**Table 5:** Table showing Root mean square error (RMSE), Max absolute error (Max abs. err.) and standard deviation (Std) for the parameter estimations of (*g_Na_, g_s_*) along the drug directions C from the experiment presented in Section 3.4.

| Sodium blocker (C) |  |  |  |
| --- | --- | --- | --- |
| Algorithms | RMSE | Max abs. err. | Std |
| Streamrak | (6.e-3, 5.e-4) | (1.e-2, 1.e-3) | (5.e-3, 5.e-4) |
| Falkon | (2.e-2, 2.e-3) | (3.e-2, 3.e-3) | (8.e-3, 4.e-4) |
| Eucl-1-nn | (3.e-2, 3.e-3) | (6.e-2, 4.e-3) | (3.e-2, 2.e-3) |
| Eucl-10-nn | (4.e-2, 5.e-3) | (6.e-2, 6.e-3) | (2.e-2, 1.e-3) |
| Apf-1-nn | (5.e-2, 2.e-2) | (1.e-1, 5.e-2) | (5.e-2, 2.e-2) |
| Apf-10-nn | (3.e-2, 1.e-2) | (5.e-2, 2.e-2) | (3.e-2, 2.e-2) |

